# Extended longevity of termite kings and queens is accompanied by extranuclear localization of telomerase in somatic organs and caste-specific expression of its isoforms

**DOI:** 10.1101/2024.03.28.587005

**Authors:** Marie Pangrácová, Jan Křivánek, Markéta Vrchotová, Hana Sehadová, Romana Hadravová, Robert Hanus, Ondřej Lukšan

## Abstract

Kings and queens of termites are endowed with an extraordinary longevity coupled with lifelong fecundity. We recently reported that termite kings and queens display a dramatically increased enzymatic activity and abundance of telomerase in their somatic organs when compared to short-lived workers and soldiers. We hypothesized that this telomerase activation may represent a non-canonical pro-longevity function, independent of its canonical role in telomere maintenance.

Here, we explore this avenue and investigate whether the presumed non-canonical role of telomerase may be due to alternative splicing of the catalytic telomerase subunit TERT and whether the subcellular localization of TERT isoforms differs among organs and castes in the termite *Prorhinotermes simplex*. We empirically confirm the expression of four in silico predicted splice variants (*psTERT1-A*, *psTERT1-B*, *psTERT2-A*, *psTERT2-B*), defined by N-terminal splicing implicating differential localizations, and C-terminal splicing giving rise to full-length and truncated isoforms. We show that the transcript proportions of the *psTERT* are caste- and tissue-specific and that the extranuclear full-length isoform *TERT1-A* is relatively enriched in the soma of neotenic kings and queens compared to their gonads and to the soma of workers. We also show that extranuclear TERT protein quantities are significantly higher in the soma of kings and queens compared to workers, namely due to the cytosolic TERT. Independently, we confirm by microscopy the extranuclear TERT localization in somatic organs. We conclude that the presumed pleiotropic action of telomerase combining the canonical nuclear role in telomere maintenance with extranuclear functions is driven by complex TERT splicing.

## INTRODUCTION

Evolution of insect eusociality was accompanied by the loss of the trade-off between fecundity and longevity that applies to solitary organisms. Reproductive castes of advanced social insects (queens of ants and eusocial bees, kings and queens of termites) are endowed with high life-long fecundity and often spectacular lifespans (Keller, 1998). Kings and queens of termites are among record holders in this respect: indirect estimates as well as empirical data suggest that they may live for more than one or even up to two decades, thus exceeding manifold the life expectancies of non-reproducing termite castes or related solitary insects (e.g., Korb & Thorne, 2017; Koubová *et al*., 2021; Monroy Kuhn *et al*., 2021). Evolutionary theories provided multiple non-exclusive arguments why should the termite kings and queens be long-lived and highly fertile at a time. Because the altruistic subfertile or sterile helpers (workers, soldiers) take over the brood care, defence and food provisioning in their colonies, the reproductives are liberated from work tasks and extrinsic mortality, which ultimately alleviates the selective pressure for precipitated reproduction in favour of delayed and long-term production of quality offspring (e.g., Keller & Genoud, 1997; Monroy Kuhn & Korb, 2016; Korb & Heinze, 2021; Tasaki *et al*., 2021; Hellemans & Hanus, 2024).

During the past few years, multiple studies addressed the proximate mechanistic aspects that allow for long life and simultaneous high fertility in termite kings and queens, converging at the statement that there is not one universal pro-longevity adaptation. Instead, multiple mechanisms are recruited by the reproductives to cope with the variety of intrinsic mortality causes. Large body of evidence has been amassed showing that termite queens and kings invest into the protection against oxidative stress, such as the antioxidant enzymes (superoxide dismutase, catalase, peroxiredoxin), vitamins and cofactors (coenzyme Q_10_, tocopherol) (Tasaki *et al*., 2017, 2019, 2023; Ye *et al*., 2019). Yet, it must be noted that the upregulation of the antioxidative mechanisms does not seem universal across termite species from different lineages, which is in line with the lack of consistent pattern of oxidative stress protection in different eusocial Hymenoptera (Parker *et al*., 2004; Corona *et al*., 2005; Kramer *et al*., 2021).

Preservation of genome integrity is another intuitive target of anti-ageing adaptations expected to occur in long-lived organisms. Indeed, mature primary queens of *Macrotermes* exhibit a markedly lower expression of transposable elements (TEs) when compared to short-lived major workers, and they probably do so thanks to significant somatic upregulation of the genes involved in the piRNA pathway, known to protect the genome integrity by TE silencing in animal germlines (Elsner *et al*., 2018; Post *et al*., 2023). This striking recruitment of germline protection in queen soma provides support to the concept of eusocial insects as superorganisms, in which the queen is analogous to the germline and workers to the soma of the superorganism (Bernadou *et al*., 2021).

Recently, termite queens (and kings) were reported to upregulate yet another mechanism of genome integrity protection, the telomerase (Koubová *et al*., 2021). Telomerase is a ribonucleoprotein with reverse transcriptase function specialized in adding short repetitive motifs to the telomeres, ends of linear chromosomes in eukaryotes. As such, it is the central element of an extremely influential paradigm stating that cell lifespan is limited by inevitable telomere shortening during mitotic divisions and that telomerase evolved to set the telomere length in germ, stem, and embryonic cells, and to compensate for telomere shortening in tissues with high lifelong proliferation. By contrast, most somatic cells of adult organisms are telomerase-negative and telomerase deregulation may lead to uncontrolled growth typical for cancer (e.g., Greider & Blackburn, 1985; Blackburn, 2005; Greenberg, 2005; Whittemore *et al*., 2019). Telomere-telomerase system has received great attention as a potentially universal lifespan regulator at cell and ultimately organismal level, with multiple evidence in humans, some other mammals and birds (Sahin & DePinho, 2010; Jaskelioff *et al*., 2011; Heidinger *et al*., 2012). Yet, research conducted across the phylogenetic diversity of animals has shown that the importance of telomere shortening and telomerase for lifespan control differ among animal lineages depending on various life history traits, such as type of postembryonic development (determinate vs. indeterminate growth), body size, general lifespan, and others (reviewed, e.g., in Smith *et al*., 2022; Čapková Frydrychová, 2023). Telomere lengths and telomerase activities may thus be efficient predictors of life expectancies in some taxa and poor in others.

In addition, multiple studies unveiled during the past two decades that telomerase may be implicated in cell lifespan and organismal fitness independently of its canonical role in telomere prolongation. Telomerase has been shown to participate in transcription regulation, DNA repair processes, apoptosis, and last, but not least, in oxidative stress protection. Some of these new functions ascribed to telomerase or its catalytic subunit TERT (telomerase reverse transcriptase) are extranuclear, especially linked with the biology of mitochondria, such as protection of mitochondrial DNA and the cell itself from reactive oxygen species, with implications for the nuclear DNA and telomere integrity (reviewed, e.g., in Saretzki, 2014; Ségal-Bendirdjian & Geli, 2019; Zheng *et al*., 2019; Smith *et al*., 2022). Non-canonical functions may also take place in cells where canonical telomerase action is of low importance, such as neurons (e.g., Eitan *et al*., 2016). The differential telomerase/TERT functions in humans has been hypothesized to be regulated by differential TERT splicing, which generates multiple TERT isoforms differing, among others, by localization signals (Avin *et al*., 2016; Ségal-Bendirdjian & Geli, 2019); dynamic evolution of TERT splicing in Metazoa is apparent from cross-taxa comparison (Lai *et al*., 2017).

There is no direct evidence that the telomere-telomerase system serves as a lifespan regulator in insects. Insect telomere and telomerase biology have historically been studied with emphasis on the diversity of oligonucleotide telomeric motifs, and presence or absence of telomerase and its activity (Frydrychová *et al*., 2004; Korandová *et al*., 2014; Kuznetsova *et al*., 2020). Rare are the studies attempting to link the life expectancies of different insects with their telomere lengths and telomerase activity. Social insects, given the great lifespan differences among castes, are an excellent model for such studies. Jemielity et al. (2007) reported shorter telomeres in short-lived males of the ant *Lasius niger* compared to the female castes, without however detecting significant differences between long-lived queens and short-lived workers. Comparison of telomere lengths in workers vs. queens of the honeybee and workers vs. kings and queens of termites converged to the same conclusions, suggesting that telomere length is unlikely to be the lifespan-limiting factor responsible for the conspicuous lifespan differences among castes in social insects (Korandová & Čapková Frydrychová, 2016; Koubová *et al*., 2021).

By contrast, striking caste-specific pattern of telomerase enzymatic activities and TERT abundances has been recorded in several social insects. It has first been reported in European honeybee, whose queens display a dramatic increase in telomerase activities in somatic organs, including those that exhibit low or no cell divisions, when compared to workers and drones (Korandová & Čapková Frydrychová, 2016; Koubová & Čapková Frydrychová, 2021). Later, we observed in several species of termites that the long-lived kings and queens show a great upregulation of telomerase activities and protein abundances in a variety of somatic organs, such as ventral nerve cord, epidermis, fat body, and others, most of which consist of post-mitotic tissues (Koubová *et al*., 2021). These results stand in contrast to telomerase activities in the solitary insect model, the American cockroach *Periplaneta americana*, characteristic by gradual decrease during the postembryonic development to low but detectable values in adult soma, a pattern typical for most vertebrates (Korandová *et al*., 2014). Hence, somatic recruitment of telomerase in termite reproductives (and honeybee queens) is yet another example of mechanisms usually proper to the germline being activated in adult social insect reproductives. Indirectly, these assumptions were confirmed in a study on bumblebees, showing that the short-lived queens of these primitive social insects lack the described pattern of somatic telomerase activation (Koubová *et al*., 2019). At the same time, presence of telomerase in organs with low or no proliferation activity strongly suggests that it may be associated with a non-canonical function in social insect reproductives, adding thus to the growing body of knowledge on telomere-independent nuclear and extranuclear pro-longevity roles of telomerase.

In this study, we build on our previous report on telomerase activation in adult soma of termite kings and queens (Koubová *et al*., 2021) and further explore the avenues defined by our observations. First, we address the question whether the presumed non-canonical role of telomerase in termites may be due to alternative splicing of *TERT* which would generate isoforms with different functions (canonical telomerase function and alternative functions) and with different subcellular locations, whose differential expression may account for observed queen- and king-specific somatic telomerase activation. Our previous work signalled the presence of up to 6 potential *TERT* isoforms in termites, and we quantify here their expression patterns across castes and organs. Second, we study the subcellular localization of TERT in different organs and castes by combining TERT protein quantifications with microscopic techniques to test the hypothesis on potential extranuclear telomerase function upregulated in king and queen soma.

In our study, we use the termite *Prorhinotermes simplex* (Rhinotermitidae), also studied as the main model in our previous paper on telomerase recruitment in the soma of reproductive castes. Our long-term observation showed that *P. simplex* primary kings and queens may live for more than 20 years in laboratory conditions (median age 10–12 years), and neotenic kings and queens regularly live for 5–10 years. By contrast, the non-reproducing helper phenotypes (workers, soldiers) rarely live more than four years (Koubová *et al*., 2021). *P. simplex* belongs among termites with linear development, whose workers retain developmental capacity to further develop into both types of reproductives (Roisin, 1988). Therefore, it is upon their differentiation into neotenic or primary reproductives (via a single nymphal stage) that the pro-longevity mechanisms are expected to be recruited, resulting in lifespan extension by up to a decade or more. Availability of multiple laboratory colonies with individually recorded age of neotenic reproductives allows obtaining sample sets with defined age of kings and queens in sufficient numbers.

## MATERIAL AND METHODS

### Termites

Multiple mature colonies of *Prorhinotermes simplex* (Rhinotermitidae; Hagen, 1858) are held at the Institute of Organic Chemistry and Biochemistry in glass vivaria lined with moistened sand (permanent dark, 27°C, 80% relative humidity). Colonies live in clusters of spruce wood slices (100×200×4 mm) and irregularly produce winged dispersers. Dozens of new colonies were established from heterosexual pairs of these dispersers between 2001 and 2014, providing thus colonies with known age of the primary reproductives. Regular inspections of such colonies allow tracking the longevity of both the primary and secondary (neotenic) reproductives, replacing the primaries after their death.

To induce the formation of neotenics, subpopulations of 50 temporary workers (pseudergates) of stages 4 or 5 (referred to as “workers” in this study) and 10 soldiers were placed in 9 cm Petri dishes lined with moistened sand and with small blocks of spruce wood offered as a diet. In these orphaned groups, multiple neotenics of both sexes start to differentiate from workers approximately 10 days after the group establishment. The groups were inspected every 72 hours, newly molted neotenics were removed, their sex scored, and they were introduced as one or two pairs into new “culture” groups of workers and soldiers, freshly removed from the original mature colony.

In experiments presented here, we used workers and neotenics of two age classes, i.e., 4–6 months old neotenics (referred to as young neotenics or young kings and queens) and mature neotenics (mature kings and queens) at least two years after their differentiation from workers.

### RNA isolation

Total RNA from termite samples was isolated using acid guanidinium thiocyanate-phenol-chloroform extraction. Briefly, samples were ground on dry ice using polypropylene pestles and homogenized in TRI reagent (Thermo Fisher Scientific). RNA was collected from aqueous phase after addition of chloroform and centrifugation according to manufacturer’s documentation, precipitated with isopropanol (1:1), washed with 80% ethanol and dissolved in nuclease-free water. RNA quantity and purity were verified using Nanodrop spectrophotometer, integrity was inspected on 1% agarose gel in TAE buffer pre-stained with GelRed nucleic acid stain (Merck).

### Next-generation Sequencing (NGS) and bioinformatic analysis

A set of total RNA isolates was prepared from heads and abdominal soma (without digestive tube) of workers and young and mature neotenic queens. In case of queens, gonads were also sampled. For each sample type, we prepared 3–4 biological replicates, each containing a pool of body parts from 3 individuals, giving rise to the total of 31 caste- and tissue-specific samples. RNA isolates were then used for high-throughput sequencing analysis (RNA-seq). Library preparation of non-stranded, poly(A)-selected cDNA libraries together with sequencing analysis on Illumina HiSeq aiming at 5 million of 2 × 150 bp paired end reads was conducted at Eurofins Genomics (Ebersberg, Germany). RNA-seq data are published in NCBI SRA database under accession numbers XYXYXYX (*to be completed upon publication*). Raw sequence reads were trimmed and filtered with the Trimmomatic v0.36 tool (Bolger *et al*., 2014) and mapped using the STAR aligner v2.7.10b (Dobin *et al*., 2013) on our unpublished in-house *P. simplex* genomic reference (see Koubová *et al*., 2021 for details on genome assembly). Reads mapping to *psTERT* coding regions were retrieved by the featureCounts v1.6.3 algorithm (Liao *et al*., 2014) and results were manually inspected. *psTERT* expression levels were calculated using common formula for FPKM values: 10^9^ × C_psTERT_ / (C_total_ × L_psTERT_), where *C* stands for read counts and *L* for gene length. Differential exon usage analysis was performed using the DEXseq package in R (Anders *et al*., 2012).

### cDNA synthesis and quantitative real-time PCR

Multiple series of RNA isolates, each originating from a different colony, were prepared from eggs (pool of 40), whole bodies of larval stages L1 (5), L2 (3) and L3 (2), workers (1), presoldiers (1) and soldiers (1). In addition, in selected colonies we prepared another series of isolates from dissected abdominal cavities of workers (2) and body cavities and gonads of mature neotenics of both sexes (1), each in at least 3 biological replicates. The cDNA was synthesized from 0.5–1 μg of RNA using the Maxima H Minus Reverse Transcriptase (Thermo Fisher Scientific) according to the manufacturer’s protocol. Isoform-specific *psTERT* expression was analyzed in a series of qPCR reactions using target-specific primers amplifying particular splice variants *TERT1, 2* (alternative splicing of exon 1), *TERT- A, B* (alternative splicing of exon 9) and glyceraldehyde 3-phosphate dehydrogenase (*GAPDH)* used as endogenous control. All primers were designed to overlap exon-junctions in order to avoid contamination with genomic DNA. Primer sequences, efficiencies and products lengths are listed in Supplementary Table S1 and Supplementary Figure S1. PCR reactions containing SYBR Green I Mastermix (Roche), primers and template in 10 μl of nuclease-free water were processed with LightCycler 480 Instrument II (Roche) as follows: 10 min initial denaturation at 95°C followed by 40 cycles of denaturation (10 s, 95°C), primer annealing (30 s, 58°C), and extension (30 s, 72°C). Reaction products were checked at the level of melting curves and verified on 2% agarose gel in TAE buffer. Relative expressions were calculated using the 2^–ΔΔCt^ method with normalization to *GAPDH* and workers as the control group.

### Protein extractions and immunochemical analysis

Protein extracts for immunochemical analyses were prepared as described in Koubová et al. (2021) with several modifications. Nuclear protein extracts were prepared as described earlier while the extranuclear protein fraction was further centrifuged at 14,000g and 4°C for 10 min to separate the cytosolic fraction and mitochondrial pellet. Mitochondrial protein was subsequently released by dissolution of membranes during 30 min incubation at 4°C in 20 mM Tris buffer, pH 7.8, containing 1.5 mM MgCl_2_, 400 mM NaCl, 20% glycerol and 1% NP-40. Protein concentrations were determined using the Micro BCA Protein Assay Kit (Thermo Fisher Scientific) according to manufacturer’s recommendations. Potential contamination among subcellular fractions was inspected by the presence of nucleus-specific marker Histone H3 on Western blot with human anti-Histone H3 antibody (ab1791) and mitochondria-specific activity of succinate dehydrogenase using the Succinate dehydrogenase activity colorimetric Assay (Sigma Aldrich), both performed according to manufacturer’s instructions.

psTERT content in somatic tissues vs. gonads and subcellular distribution in extranuclear and nuclear space was inspected in a series of direct enzyme-linked immunosorbent assays (ELISAs). Protein extracts were obtained from dissected abdominal cavities without digestive tube of 2 pooled individuals in 3–4 replicates for each studied phenotype: workers, and neotenic kings and queens of two maturation stages. In case of neotenics, we also prepared protein extracts from reproductive organs. We applied 2 µg of protein extract per well in 3 technical replicates followed by incubation overnight, blocking in 3% bovine serum albumin (BSA), incubation with primary antibody anti-psTERT (1:1000; for details on antibody origin see Koubová *et al*., 2021), secondary antibody horseradish peroxidase (HRP)-conjugated Goat Anti-Rabbit IgG (1:5000; Abcam), both in blocking solution and detection using the Supersensitive TMB substrate (3,3’,5,5’-tetramethylbenzidine, Sigma-Aldrich) according to the manufacturer’s instructions. All steps were interleaved by 3 washes with Tris-buffered saline (TBS) with 0.1% Tween 20, the last washing step preceding TMB application was carried out with TBS without detergent.

Cytosolic, nuclear and mitochondrial protein fractions for Western blot analyses were obtained from whole bodies without digestive tract using a pool of 40 workers, quantified and precipitated with 10% ice-cold trichloroacetic acid followed by two washing steps with 100% ice-cold acetone. Dried pellet was dissolved in modified Laemmli sample buffer (125 mM Tris-base, pH 6.8, 0.5% mercaptoethanol, 2.5% SDS, 50 mM NaOH, 10% glycerol and 0.006% bromophenol blue) and boiled for 10 min. Proteins were separated using Sodium dodecyl sulfate polyacrylamide gel electrophoresis (SDS PAGE) in 8% polyacrylamide gel together with PageRuler protein ladder (Thermo Fisher Scientific) and transferred onto polyvinylidene difluoride membrane (PVDF). The membranes were then blocked using 3% nonfat dried milk in TBS with 0.1% Tween 20 and incubated with antibodies described above. Immunoreactive bands were visualized by detection of chemiluminescent signals after addition of the Pierce ECL detection system (Thermo Fisher Scientific). All ELISA and Western blot analyses were confirmed in at least 3 independent experiments with individuals originating from different colonies.

### Cell proliferation mapping using 5-ethynyl-2’-deoxyuridine (EdU)

To track the cell proliferation rate in different organs of *P. simplex* castes, we used the Click-iT EdU Alexa Fluor 488 Imaging Kit according to the manufacturer’s protocol (Invitrogen by Thermo Fisher Scientific). Cold-anesthetized termites (5 min at 4°C) were injected with 100 nl of 40 mM EdU in distilled water into the thorax by the hind leg articulation using a 5 ml Hamilton microsyringe with a custom Hamilton needle (34/17/pstN/tapN)S (Chromservis, Prague, Czech Republic). Injected individuals were marked and placed into Petri dishes with groups of untreated workers and soldiers and sampled after 1, 2, 3, and 7 days.

Dissected organs were fixed with 4% formaldehyde in phosphate buffer (PB) for 2 hours and then washed two times in PB and once in PB supplemented with 0.5% Triton X 100 (PB-T) for 15 min each at room temperature (RT). Samples were blocked with 3% BSA in PB for 30 min at RT, incorporated EdU was labeled by Alexa Fluor azide in a Click-iT reaction overnight at 4°C according to manufacturer’s instructions. After rinsing with PB (three times for 15 min at RT), the samples were treated with 1 µg/ml 4’-6-diamidino-2-phenylindole (DAPI) for 10 min at RT. After stopping the reaction in distilled water, the samples were dehydrated in an ethanol series (50%, 70%, 90%, and 100% for 15 min each) and mounted into methyl-salicylate. Samples were visualized under laser scanning confocal microscope FLUOVIEW FV1000 (Olympus, Tokyo, Japan). Two to three individuals were observed from each studied caste (workers, presoldiers, soldiers and mature neotenics of both sexes).

### Immunohistochemical detection and microscopy

Cold-anesthetized animals were penetrated by tungsten needle in sterile saline and submerged in a fixative of saturated picric acid, 4% formaldehyde, and 2.3% copper acetate supplemented with mercuric chloride (Bouin-Hollande solution, Levine et al. 1995) overnight at 4°C. The fixative was then thoroughly washed with 70% ethanol. Standard techniques were used for tissue dehydration, embedding in paraplast, sectioning to 7 µm, deparaffinisation and rehydration. The sections were treated with Lugol’s iodine followed by 7.5% solution of sodium thiosulphate to remove residual heavy metal ions, and then washed in distilled water and phosphate-buffered saline supplemented with 0.3% Tween 20 (PBS-T). The nonspecific binding sites were blocked with 5% normal goat serum in PBS-T (blocking solution) for 30 min at RT. Incubation with anti-psTERT primary antibody diluted 1:200 in the blocking solution was performed in a humidified chamber overnight at 4°C. Pre-immune serum was used instead of the primary antibody in the control staining. After a thorough rinsing with PBS-T (three times for 10 min at RT), the sections were incubated in the goat anti-rabbit IgG conjugated Alexa Fluor 488 (Life Technologies, Carlsbad, CA, USA) diluted at 1:500 in the blocking solution for 1 h at RT of and further rinsed with PBS-T (three times for 10 min at RT in the dark). Stained sections were dehydrated, mounted in DPX mounting medium (Fluka, Buchs, Switzerland)), and studied under the laser scanning confocal microscope FLUOVIEW FV1000 (Olympus, Tokyo, Japan). Two to three individuals were observed from each studied caste (workers, presoldiers, soldiers and mature neotenics of both sexes).

### Immunogold labeling and transmission electron microscopy (TEM)

Whole animals or dissected organs were fixed for 2h at 4°C in a solution consisting of 2% formaldehyde in PBS pH 7.4. After washing the samples were gradually dehydrated at 4°C in 30%, 50%, 70%, 90% aqueous ethanol. After dehydration the samples were embedded in LR White embedding medium. Polymerization was carried out at 4°C under UV light for 72hr.

Parlodion-coated nickel grids carrying ultrathin sections (70 nm) were washed briefly in distilled water and blocked for 1 h with blocking buffer consisting of 10% normal goat serum (v/v), 1% BSA (w/v) in PBS (pH 7.4). Then the grids were incubated o/n at 4°C with the anti- psTERT antibody diluted 1:100 in PBS pH 7.4 containing 0.5% BSA (w/v) and 0.05% (v/v) Tween 20 (buffer A). After incubation the grids were three times washed in PBS (pH 7.4) with 0.1% BSA (w/v) and then transferred to a droplet of goat anti-rabbit IgG conjugated to 10-nm gold particles (British Biocell) diluted 1:25 in buffer A and incubated 1 h at 4°C. After washing, the grids were finally stained with uranyl acetate and examined with a JEM 2100 Plus electron microscope (JEOL, Japan).

### Statistics

Differential exon usage analysis was performed using the DEXseq statistic package (Bioconductor). Caste- and tissue-specific *psTERT* expression was compared using RNA-seq FPKM values and two-way ANOVA with caste and tissue set as factors, expression in gonads of young and mature neotenic queens was compared using two-tailed Student’s t-test.

Data from isoform-specific expression analyses and from ELISA experiments were related to the values recorded in workers (except for ratios of isoforms and ratios of extranuclear and intranuclear TERT) and the means then compared using one-way ANOVA, followed either by Dunnett’s post hoc test with worker values set as controls or by Bonferroni-corrected multiple comparisons between pairs of relevant phenotypes (castes, organs, isoforms). Data from ELISA quantification in cell compartments of workers and queens were tested using two-tailed Student’s t-test in each compartment. Prior to parametric statistics listed above, the datasets were tested for equality of variances (Brown-Forsythe test), and in some cases log_2_ transformed to reduce their heteroscedasticity and meet the requirements for parametric analysis. Tests and data transformations used are indicated in figure legends of each graph, basic statistics of all tests are listed in Electronic Supplementary Material.

## RESULTS

### *psTERT* and alternative transcript variants

*psTERT* is expressed in multiple isoforms due to three alternative transcription start sites and two alternative splicing events localized in exons 1 (*TERT1* and *2*, also referred to as N-terminal isoforms) and 9 (*TERT-A* and *B*, C-terminal isoforms) (Figure 1A). All three N-terminal protein isoforms are in frame and differ only in the leading sequences – a putative signal peptide was predicted *in silico* in *TERT1*, while *TERT2* carries a strong monopartite nuclear signal RRRKKKIK in the proximal N-terminal part. The *TERT3* isoform is transcribed from an alternative start site located in exon 3 and has exactly the same sequence as the other transcript variants. It is barely recognizable in the background of the other isoforms and has no obvious functional features, and we therefore decided to exclude this variant from further analyses. The splice variants *TERT-A* and *TERT-B* differ in the C-terminal extension, with the truncated isoform *TERT-B* lacking conserved motifs (E-III, E-IV) that have been reported to have functional impact on the full activity of the TERT holoenzyme in humans (Banik *et al*., 2002). Hence, we further studied the *psTERT* expression at the gene level covering all possible isoforms, as well as at the level of individual exons with emphasis on potential caste-specific expression of N- and C-terminal splice variants.

**Figure 1.**
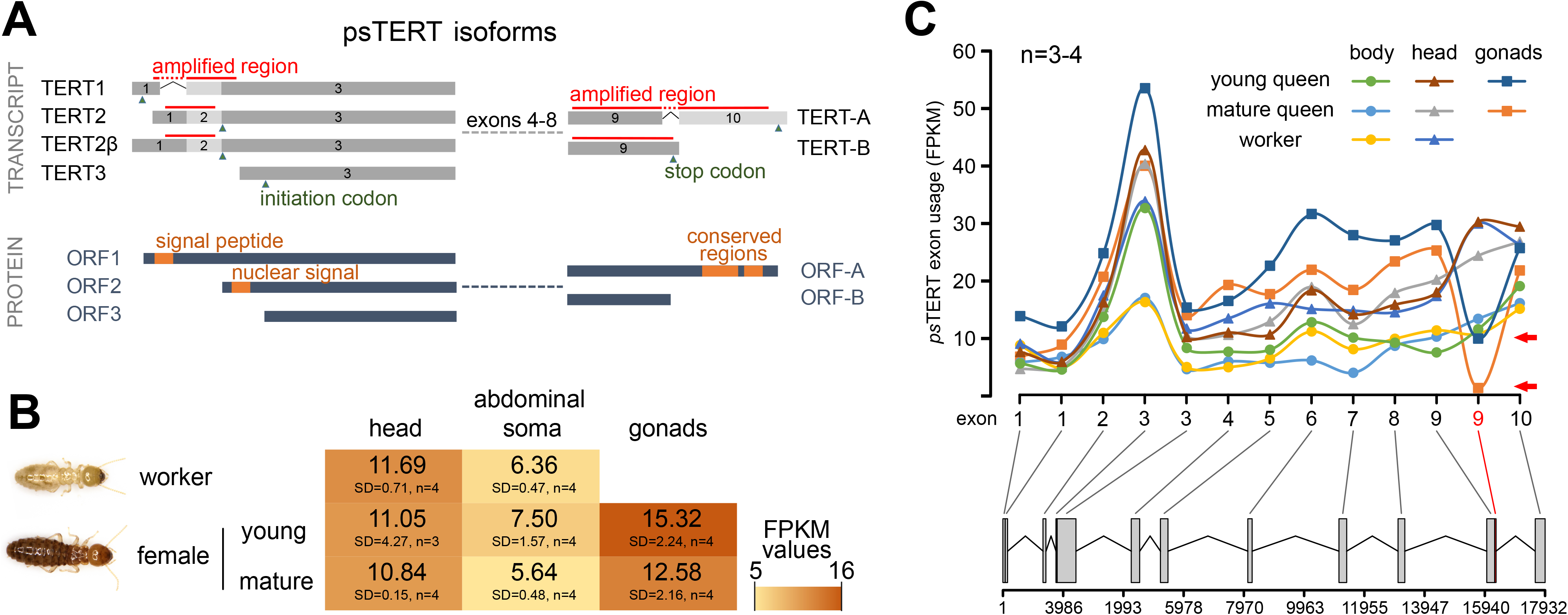
*psTERT* gene and protein structure, expression at gene and exon levels in workers and queens. **A.** Transcript and protein structure of *psTERT* isoforms with emphasis on alternative splicing in exons 1 and 9. Regions used for qPCR amplification and isoform-specific expression analysis are marked by red bars, initiation and termination codons by green triangles and domains related to telomerase localization or full activity (putative signal peptide, nuclear signal and C-terminal conserved regions) are highlighted as orange boxes. **B.** Heatmap representing normalized FPKM levels of *psTERT* expression in multiple body parts of *P. simplex* workers and young (<3 months) and mature (>2 years) neotenic queens. **C.** NGS-based analysis of differential exon usage in head, abdominal somatic organs and gonads of workers and young and mature neotenic queens. The plot represents means of FPKM normalized counts for each exonic part calculated from from 3-4 replicates, statistical analysis was performed using the DEXseq package (Bioconductor). Lower part of the plot represents gene structure, gene regions with significantly differential exon usage are plotted in red and marked with red arrow.

### *psTERT* expression at gene and exon level

Figure 1B shows a comparison of *psTERT* transcript abundances at the gene level covering all possible isoforms in the abdominal soma and head of workers and neotenic queens at two maturation stages (and in the queen gonads) using low-coverage RNA-seq. *psTERT* expression ranked as gonads > head > abdominal soma. While there were significant differences between heads and abdominal soma, no flagrant differences were observed for these body parts between workers and the maturation stages of queens (Figure 1B, Supplementary Table S2) (caste effect, F=2.13, p=0.15; body part effect, F=24.3, p=10^-4^; interaction, F=0.09, p=0.92). Likewise, there was no significant difference in *psTERT* transcript abundance between the gonads of young and mature queens (t=1.496, p=0.19).

In the next step, we used the same data and the Bioconductor DEXseq package to assess differences at the *psTERT* exon level, which could indicate caste- or tissue-specific expression of different *psTERT* splice variants. Utilization of all exons was highly consistent in all groups except for 3’-terminal part of exon 9, which is spliced into transcript variants containing exon 10 and results in translation of the full-length psTERT open reading frame (ORF). The terminal part of exon 9 was significantly underrepresented in the reproductive organs of both, young and mature neotenic queens compared to the worker and queen soma (p<10^-3^). Differential exon usage is demonstrated by a significant decrease in mean FPKM in Figure 1C. These observations indicate that different psTERT isoforms are present in gonads and somatic organs.

### Expression of *psTERT* splice variants

In a series of PCR reactions, we first examined the presence of all potential N- and C-terminal splice variants listed above and in Figure 1A in cDNA prepared from heads and abdominal soma of *P. simplex* soldiers by combining forward primers specific for *TERT1* and *2* isoforms with reverse primers specific for *TERT-A* and *B*. We found that all four combinations *TERT1-A, TERT1-B, TERT2-A,* and *TERT2-B* are co-expressed in both tissues (Supplementary Figure S1, S2, Supplementary Table S1). In addition, we discovered a novel isoform *TERT2*β that contains a complete and unspliced exon 1 resulting in the same ORF as in the case of previously characterized *TERT2* (Figure 1A).

We next focused on the expression of specific C- and N-terminal splice variants at different developmental stages and in somatic tissues compared to the germline in adult neotenic kings and queens. Guided by the previous observations in the NGS data, we first examined the expression of isoforms that differ in splicing of exon 9. In the first experiment, we studied *TERT-A* and *B* levels during development from embryo through the three larval stages L1–3 to worker and the transition from worker through the presoldier stage to fully developed soldier (Figure 2A, B, Supplementary Table S3a). Both isoforms were the most abundant in eggs (four times larger expression compared to workers, p<10^-4^), and their expression gradually decreased through the larval development to workers to reach the lowest values in soldiers (in case of *TERT-A* significantly lower than in workers, p<10^-2^). The relative ratio of both variants (*TERT-B*:*TERT-A*) did not differ dramatically in the examined phenotypes compared to workers, only a slight preference for the full-length *TERT-A* in embryonic cells and for the truncated *TERT-B* in soldiers was observed (both with p<0.05; Figure 2C, Supplementary Table S3a).

**Figure 2.**
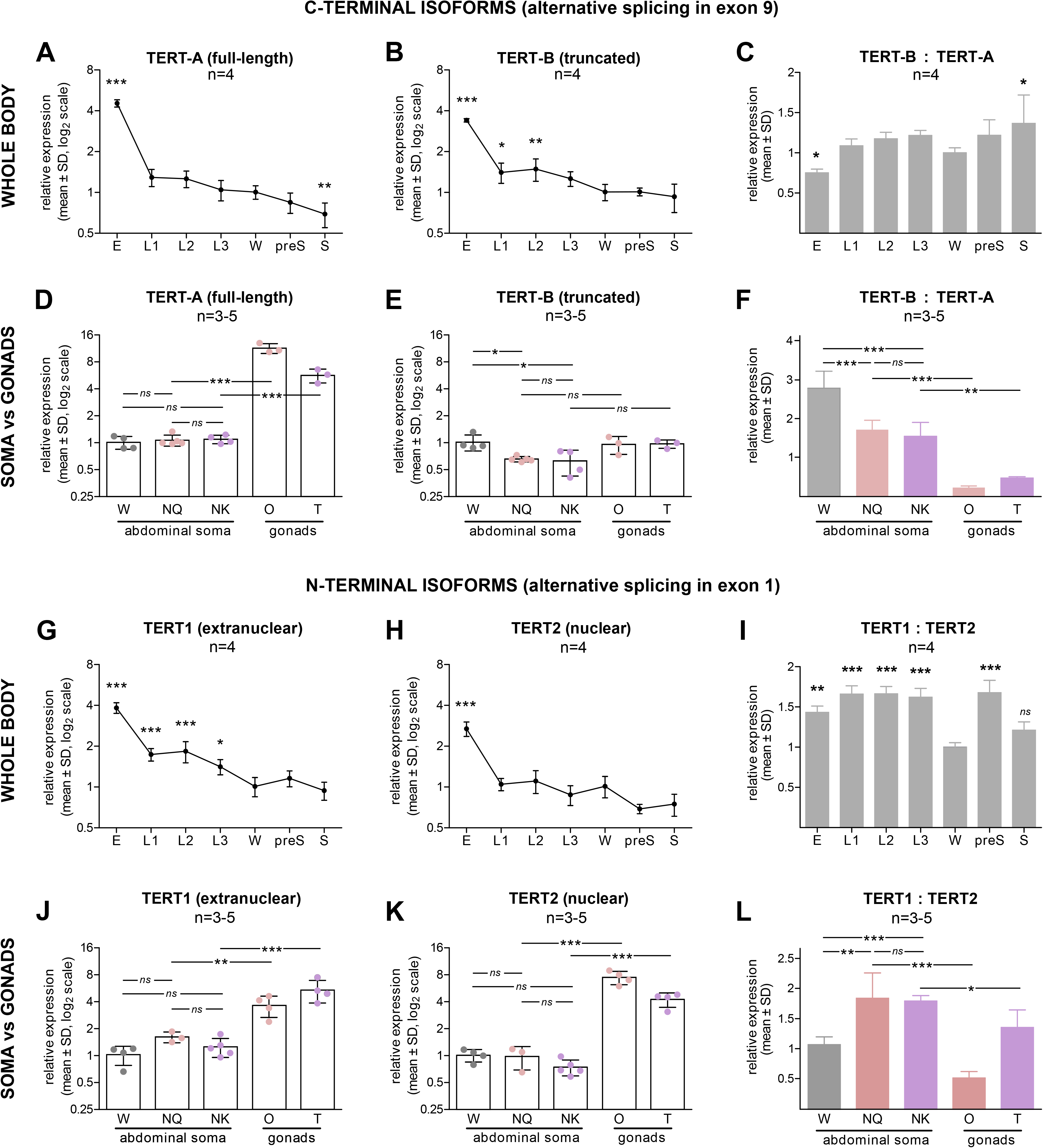
Relative isoform-specific *psTERT* expression across *P. simplex* ontogeny and in workers vs. neotenic reproductives. Left and middle graphs show expression levels of the C-terminal (A, B, D, E) and N-terminal isoforms (G, H, J, K) related to the worker values. Right graphs (C, F, I, L) represent their relative ratios. **A-F**. Expression levels of alternatively spliced C-terminal isoforms psTERT-A and B in whole body RNA extracts of multiple life stages (A-C) and in abdominal soma of workers and soma and reproductive organs of neotenics (D-F). **G-L**. Expression levels of alternatively spliced N-terminal isoforms psTERT1 and 2 in whole body extracts of multiple life stages (G-I) and in abdominal soma of workers and soma and reproductive organs of neotenics (J-L). The data in graphs A, B, D, E, G, H, J, K are related to the average value in the abdominal soma of workers from the respective colonies, log_2_ transformed and compared using one-way ANOVA followed by Dunnett’s post hoc test with worker values set as control (A, B, G, H) or by Bonferroni- corrected multiple comparison (D, E, J, K). The data in graphs C, F, I, L represent the ratio of non-transformed transcript abundances for two different isoforms, either previously related to the worker value (C, I) or not (F, L), compared using one-way ANOVA followed by Dunnett’s post hoc test with worker values set as control (C, I) or by Bonferroni-corrected multiple comparison (F, L). P-values reported as *, p < 0.05; **, p < 0.01; ***, p < 0.001. Detailed test statistics are listed in Supplementary Table S3.

To further explore the results of the exon usage analysis, we then decided to set up another experiment comparing the expression of both C-terminal isoforms in the abdominal soma of workers and mature neotenic kings and queens, and in the reproductive organs of neotenics. As shown in Figure 2D and Supplementary Table S3a, there were no significant differences between somatic transcript levels in workers and neotenics for the full-length *TERT-A* isoform, but dramatically higher levels of *TERT-A* were observed in the gonads of kings and queens compared to their somatic tissues (p<10^-3^).

The situation was different for the truncated *TERT-B*, which was slightly but significantly underrepresented in the soma of reproductives compared to workers (p<0.05), while no noticeable differences between soma and gonads were found in neotenics. The differences were conspicuous when relative ratios of *TERT-B*:*TERT-A* were compared; the full-length *TERT-A* was highly significantly (p<10^-3^) enriched in the soma of kings and queens (*TERT-B*:*TERT-A* ≈ 1.6:1) compared to workers (≈ 2.8:1) and in the reproductive organs of neotenics (0.23:1 in female gonads, 0.47:1 in male gonads) compared to their soma (p<10^-2^ for kings and 10^-3^ for queens, Figure 2F, Supplementary Table S3a).

Taken together, these observations reveal two significant trends for the C-terminal isoforms. First, the abundance of both isoforms decreases during the development of an egg through the larval stages to the mature sterile phenotypes (workers and soldiers), and the ratio of the full-length *TERT-A* isoform to the truncated *TERT-B* isoform was highest in eggs and tended to decrease during post-embryonic development. Second, somatic transcript abundance of the truncated *TERT-B* isoform is higher in workers than in neotenic kings and queens, and the full-length *TERT-A* variant is dramatically overrepresented in the gonads of neotenics, leading to highly significant differences in the representation of the full-length isoform compared to the truncated isoform in the ranking of gonads ≫ soma of kings and queens ≫ soma of workers (Figure 2F, Supplementary Table S3a).

Next, we applied the same workflow to N-terminal splice variants. We again observed a significant overexpression of both variants, *TERT1* (with predicted signal peptide suggesting extranuclear localization) and *TERT2* (with predicted nuclear localization) in RNA isolates from eggs, indicating activation of telomerase at the level of both isoforms in the embryo (p<10^-3^), followed by a decrease in their abundance along the development of sterile castes (workers and soldiers) (Figure 2G, H, Supplementary Table S3b). The ratio between *TERT1* and *TERT2* was significantly higher in all larval stages, presoldiers (both p<10^-3^) and embryos (p<10^-2^) compared to workers, while there was no significant difference when comparing soldiers and workers (Figure 2I, Supplementary Table S3b).

The somatic transcript abundances of *TERT1* and *TERT2* did not differ significantly among workers, kings and queens, but both isoforms were highly significantly upregulated in the gonads compared to the soma (p<10^-2^ and 10^-3^ in queens, 10^-3^ in kings) (Figure 2J, K, Supplementary Table S3b). Examination of transcript abundance as the ratio between extranuclear *TERT1* and nuclear *TERT2* revealed highly significant differences between the soma of workers (*TERT1*:*TERT2* ≈ 1:1) and that of both sexes of reproductives (≈ 1.8:1, p<10^-2^ for queens and 10^-3^ for kings). At the same time, the ratio was significantly higher in the soma of neotenics than in their gonads (≈ 0.5:1 for queens, p<10^-3^; ≈ 1.3:1 p<10^-1^ for kings) (Figure 2L, Supplementary Table S3b).

To summarize, the N-terminal variants showed two significant trends. First, their transcript abundances decreased during the development from eggs through larval stages to workers and soldiers, and the relative abundances of the putative extranuclear isoform were higher in larval stages compared to workers and soldiers. Second, the nuclear isoform was more abundant in the gonads of neotenics than in their soma, and the ratio of extranuclear to nuclear isoform was higher in the soma of kings and queens than in that of workers and in gonads.

### psTERT protein abundance and its subcellular distribution

Our previous work reported a significant and age-dependent increase in psTERT protein abundance in somatic tissues of reproductives compared to workers (Koubová *et al*., 2021). Here, we decided to build on these observations by quantifying TERT abundances in different cell compartments of somatic tissues using ELISA experiments with affinity-purified rabbit polyclonal antibody against psTERT prepared using recombinant protein corresponding to the C-terminal part of psTERT.

First, we deciphered TERT abundance in nuclear and extranuclear protein fractions from the abdominal soma of workers and two maturation stages of neotenic kings and queens (4–6 months old, >2 years old). As shown in Figure 3A and Supplementary Table S4, extranuclear TERT content was significantly higher in both maturation stages of queens (p<10^-1^ for young queens, p<10^-2^ for mature queens) and in young kings (p<10^-2^) compared to workers, while the higher amount of TERT in mature kings was found to be non-significant. In contrast, TERT abundance in the nuclear fraction was found equal or non-significantly lower in reproductives compared to workers. When the values obtained were expressed as a ratio of extranuclear to nuclear somatic TERT, the trend of extranuclear increase in reproductives was even more convincing, as the ratio was significantly to highly significantly greater in all neotenic phenotypes compared to workers (Figure 3B, Supplementary Table S4).

**Figure 3.**
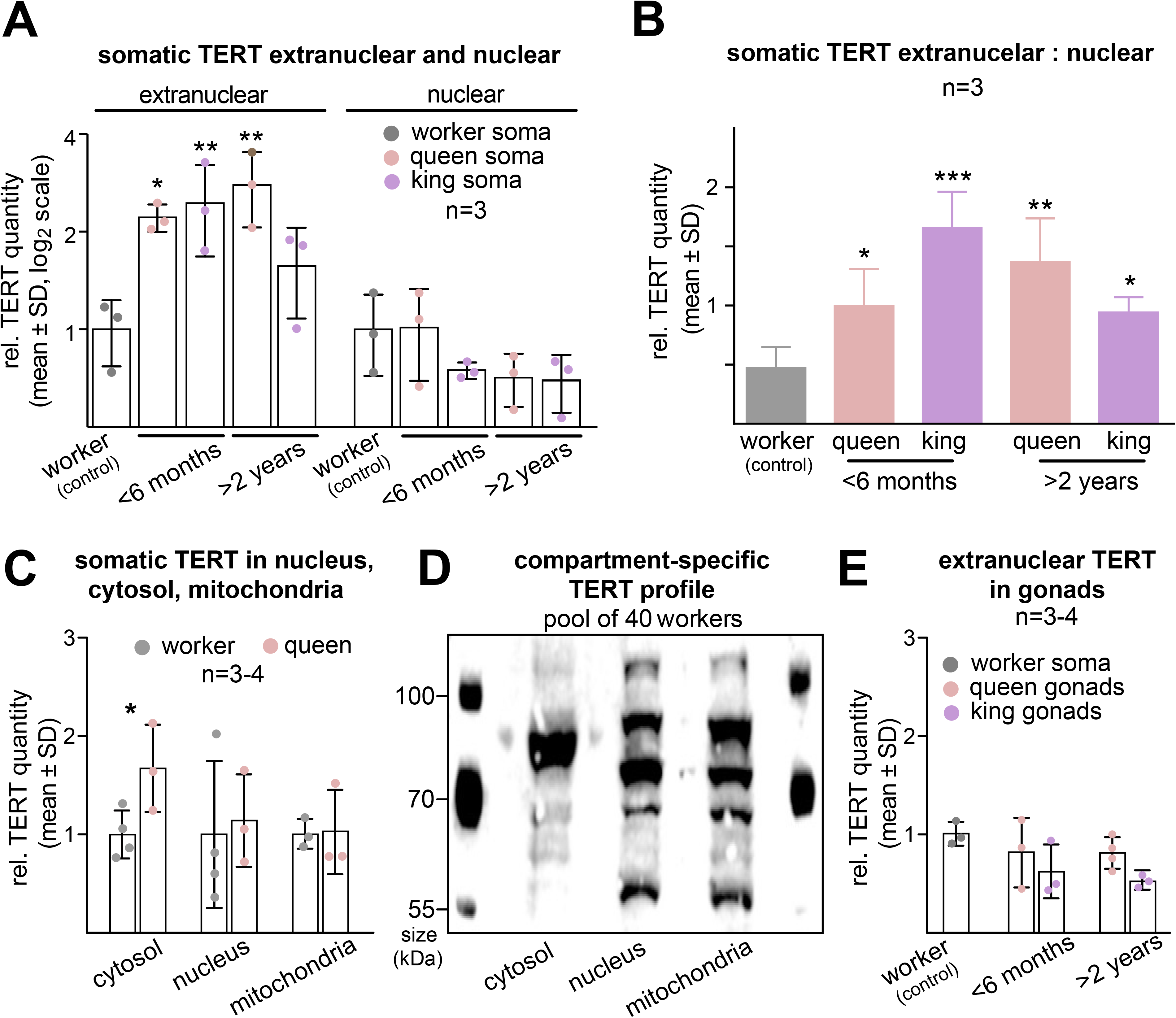
psTERT protein abundances and subcellular distribution. **A.** Relative quantity of extranuclear and nuclear psTERT in abdominal soma of workers and neotenic kings and queens of two maturation stages (<6 months, >2 years) related to the values in workers. **B.** Ratio of extranuclear to nuclear quantities of psTERT in abdominal soma of workers and neotenic kings and queens of two maturation stages (<6 months, >2 years), without previous division by worker values. **C.** Relative quantity of somatic psTERT in cytosolic, nuclear and mitochondrial protein fraction of workers and young (<6 months) neotenic queens related to the values in workers. **D.** Differential representation of psTERT isoforms in cytosolic, nuclear and mitochondrial protein extracts from 40 *P. simplex* workers visualized using Western blot. **E.** Relative quantity of extranuclear psTERT in worker soma and in gonads of neotenic kings and queens of two maturation stages (<6 months, >2 years) related to the values in workers. In A, B and E, the means were compared using one-way ANOVA followed by Dunnett’s multiple comparison with workers values set as a control, in A and E the data was log_2_-transformed prior to analysis. In C, the means in workers and queens were compared independently for each compartment using a Student’s t-test. P-values reported as *, p < 0.05; **, p < 0.01; ***, p < 0.001. Detailed test statistics are listed in Supplementary Table S4.

Next, we compared psTERT abundance in three cellular compartments, i.e., the nuclear, cytosolic and mitochondrial fractions of the abdominal soma of workers and 4–6-month-old neotenic queens. As shown in Figure 3C, we only observed a significant increase in cytosolic TERT content of queens compared to workers, while the nuclear and mitochondrial fractions did not show significant differences (Supplementary Table S4). The observed caste-specific psTERT dynamics in the cytosolic, but not in the nuclear and mitochondrial fractions prompted us to compare the patterns of TERT isoforms in the three compartments using Western blot with high-concentration protein extracts from a pool of 40 workers. The analysis revealed a clearly different pattern of psTERT isoforms ranging between 60 and 130 kDa in the cytosolic fraction compared to the mitochondria and nucleus where the profiles were highly similar (Figure 3D, Supplementary Table S4).

Since all these results pointed at the caste-specific increase in somatic cytosolic TERT in *P. simplex* neotenics, we decided to evaluate the dynamics in extranuclear psTERT in protein extracts from gonads of neotenics of both sexes and two age classes. Unlike in somatic organs, extranuclear psTERT abundance in gonads showed no significant dynamics in neotenics and remained lower than that in control somatic extranuclear extracts from workers (Figure 3E, Supplementary Table S4).

### Cellular localization of psTERT

Independently of the experiments reported above, we addressed the question of cellular psTERT localization in somatic tissues of *P. simplex* castes using immunohistochemical analyses with anti-psTERT antibodies. In particular, we focused on abdominal organs that show the highest upregulation of telomerase activity in reproductives and where canonical telomerase activity was least expected due to low cell division rates in the late stages of post-embryonic development, i.e. the ventral nerve cord and fat body (Koubová *et al*., 2021). Indeed, no dividing cells could be observed in the ventral nerve cord of workers and mature neotenics of both sexes in EdU incorporation experiments within 72 hours, whereas extensive cell proliferation occurred in the midgut section of the digestive tube (Figure 4A, B, E, F, I, J). In all three castes examined, the sections of abdominal ganglions revealed a clear extranuclear anti-psTERT signal in the neuronal bodies (Figure 4C, D, G, H, K, L), although the quantitative differences between workers and reproductives previously observed for telomerase activities were not evident. In abdominal sections of fat bodies, only a few single cell divisions occurred within 72 hours in workers, neotenic kings and queens (Figure 4M, P, S). In most parts of the fat body, anti-psTERT fluorescence was observed as scattered extranuclear signals in the cytoplasm surrounding the large lipid droplets that dominated the adipocyte cells (Figure 4O, R, U, Supplementary Figures S3, S4 and S5); only in a minor portion of cells a weak nuclear signal was also detected, as seen in the samples of mature kings in Figure 4U. Quantitative differences between workers and reproductives could not be inferred from comparison of psTERT signals. Detailed observations of anti-psTERT signals in both, the sections and whole mount preparations of workers, suggested the extranuclear localization of psTERT also in the neuronal bodies of the brain (Supplementary Figure S3).

**Figure 4.**
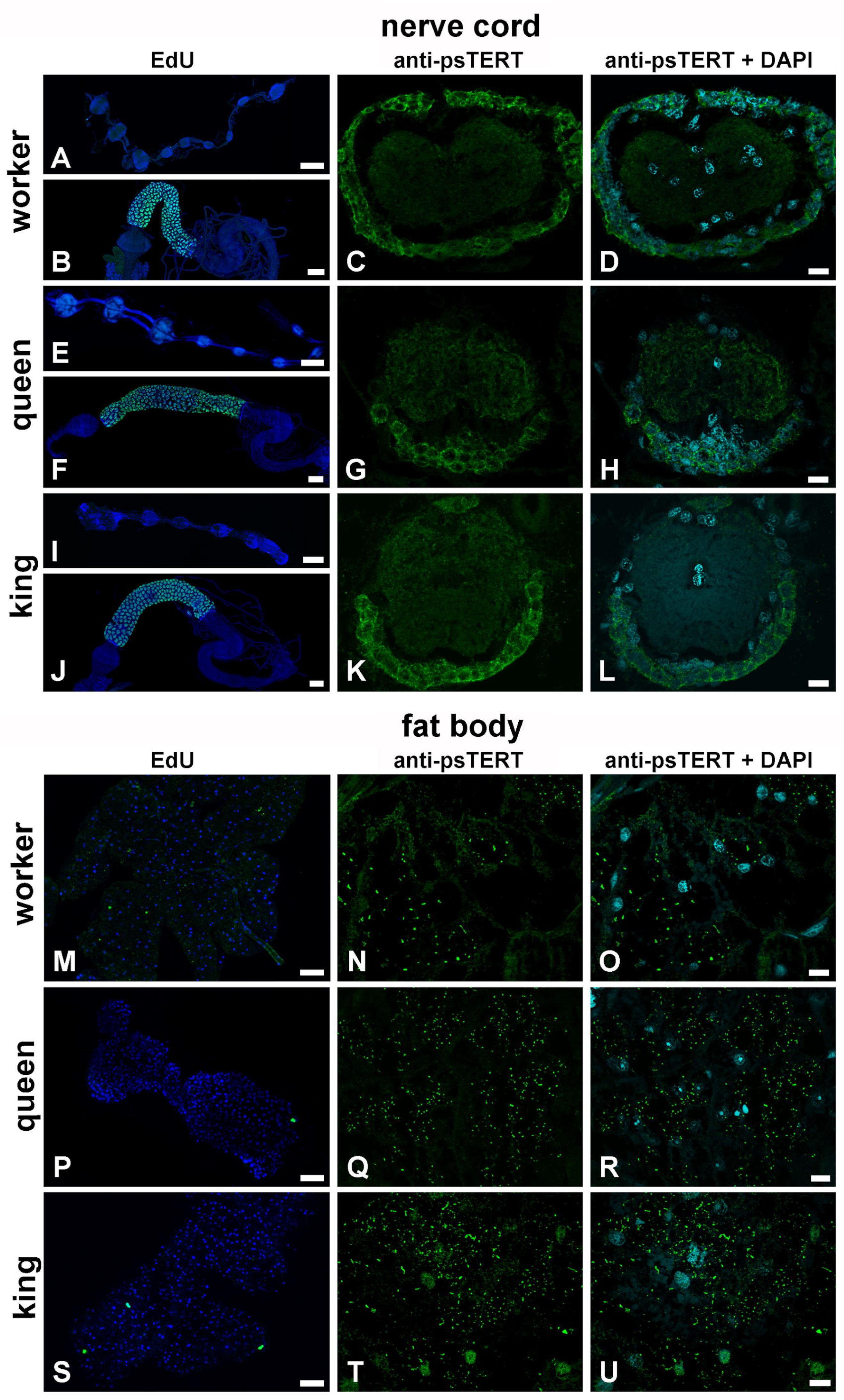
Immunodetection of DNA synthesis and psTERT in somatic tissues of P. simplex. **A-L**. Localization of EdU incorporated during active DNA synthesis within 72 h show no evidence for nuclear divisions in the whole mount preparations of nerve cord from workers (A), mature neotenic queens (E), and kings (I) but intense DNA synthesis in the control samples of the midgut from workers (B), mature neotenic queens (F), and kings (J), where the green-blue color represents colocalization of EdU signals (green) and DNA labelled by DAPI (blue). Anti-ps-TERT immunoreactivity (green) is presented separately and merged with DAPI staining (blue) in cross sections of an abdominal neuronal ganglion in workers (C, D), mature neotenic queens (G, H) and kings (K, L). **M-U**. The green color in first column indicates localization of EdU incorporated in active DNA synthesis in the nuclei of the abdominal fat body of workers (M), mature neotenic queens (P) and kings (S) during 72 h and the blue color shows DNA labelled by DAPI. Immunolabelling with anti-ps-TERT antibody (green) is shown separately and merged with DAPI staining (blue) in the sections of abdominal fat body in workers (N, O), mature neotenic queens (Q, R) and kings (T, U). Scale bars indicate 100 µm in whole mount preparations and 10 µm in tissue sections.

In parallel with the immunohistochemical analyses on paraffin sections and wholemount tissues, we applied immunogold anti-psTERT labeling to ultrathin sections from the abdominal cavities of three mature neotenic *P. simplex* queens and to dissected queen abdominal organs, including the ventral nerve cord, fat body, gonads, and muscle tissues, and visualized them using TEM. In the sections of ventral nerve cord ganglion, we detected a positive psTERT signal distributed across the neuronal cytoplasm and nuclei (Supplementary Figures S5E, S6C). Extranuclear localization associated with distribution in the mitochondria was observed in some sections of the fat body (Supplementary Figure S6E), while most of the fat body signal was localized in cytoplasmic granular structures most likely represented by protein granules (Figure 5B, Supplementary Figures S6K, S7). psTERT was also detected in the cytosol and mitochondria in muscles (Supplementary Figure S6O) and in nuclei and mitochondria of primary oocytes in ovaries (Supplementary Figure S6S). No signal was observed in control sections processed without primary antibody (Figure 5C, F and Supplementary Figures S6D, H, L, P, T).

**Figure 5.**
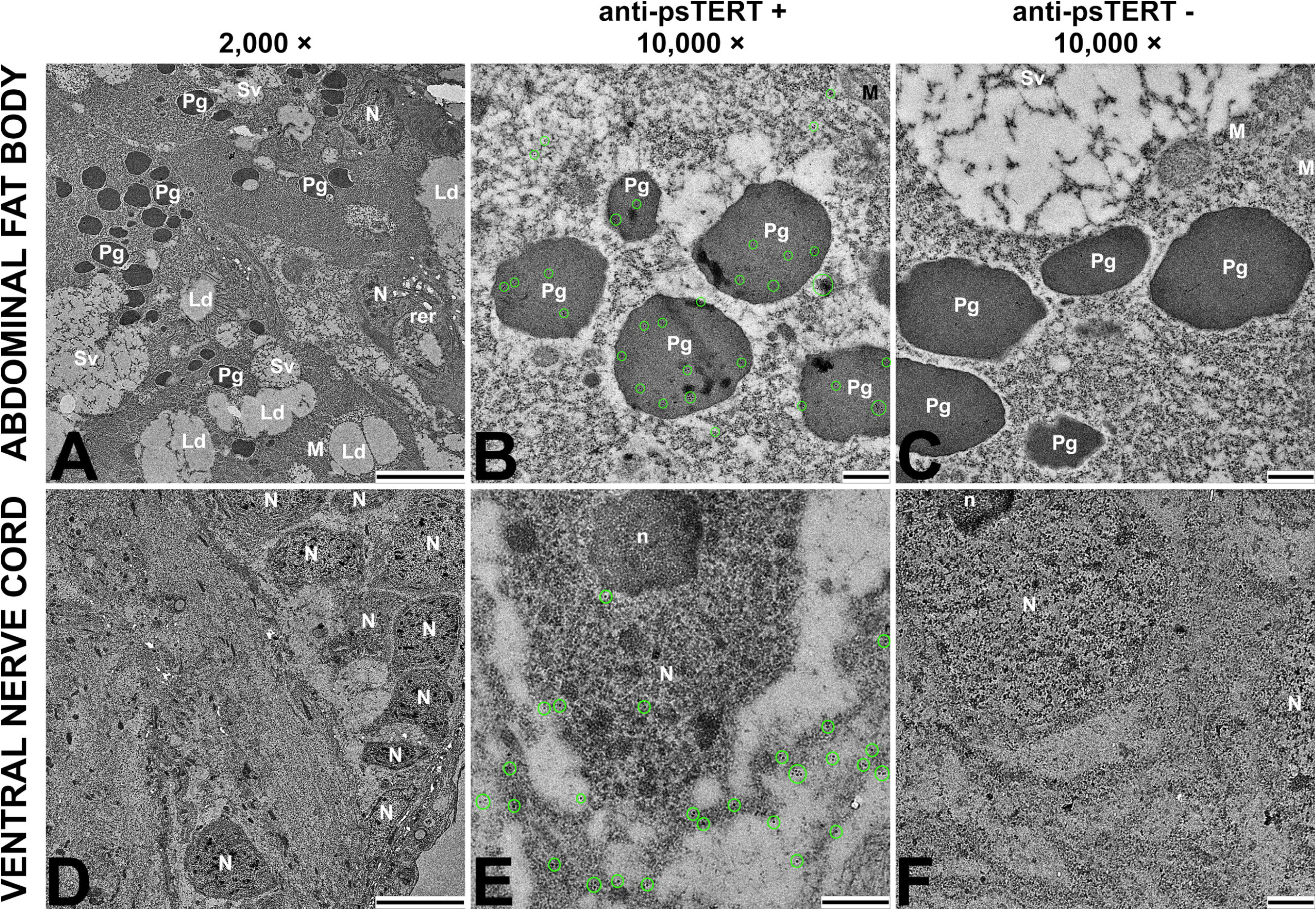
Transmission electron microscopy (TEM) with psTERT detection using anti- psTERT antibody and immunogold labeling. Immunoreactions were performed on ultrathin sections of fat body (A–C) and ventral nerve cord (D–F) from mature *P. simplex* neotenic queens. **A, D.** TEM micrographs at 2,000× magnification showing structure of examined tissues, scale bar 5 μm. **B, E.** TEM micrographs at 10,000× magnification with immunochemical detection of psTERT, scale bar 500 nm. Positive signal represented by black dots with white margin is highlighted with green circles. **C, F**. TEM micrographs at 10,000× magnification from control samples processed without primary antibody, scale bar 500 nm. Cellular organelles are marked with white symbols: N - nucleus, n - nucleolus, M - mitochondria, Ld - lipid droplets, Sv - secretory vesicles, Pg - protein granules, rer - rough endoplasmic reticulum.

## DISCUSSION

For a long time, it was assumed that telomerase reverse transcriptase is only active in the cell nucleus as part of the telomerase holoenzyme, which is responsible for protecting telomeres from wear and mitosis-dependent shortening. However, the growing list of alternative nuclear and extranuclear functions attributed to telomerase/TERT calls for new perspectives on this pleiotropic enzyme. Recent research in honeybees and termites suggests that telomerase may also play non-canonical roles in these eusocial insects, which appears to be related to the exceptional longevity of kings and queens (Korandová & Čapková Frydrychová, 2016; Koubová *et al*., 2021), reviewed in (Čapková Frydrychová, 2023). Our previous research in several termite species has shown that their long-lived reproductive castes have high enzymatic activities of telomerase and TERT abundance in their somatic tissues, including post-mitotic organs. This observation was in contrast to the general expectation of telomerase down-regulation in adult phenotypes known from most vertebrates, as well as with the data on solitary insects and non-reproducing termite phenotypes, where telomerase activities are detectable but low (Korandová *et al*., 2014; Koubová *et al*., 2021). These results pointed at the potential telomere-independent pro-longevity function of telomerase/TERT in termites and prompted the two questions addressed in the present follow-up study. First, is there an indication that differential splicing of TERT may be responsible for the distinction between the canonical role of telomerase best demonstrated in the germline, and the non-canonical role that presumably occurs in the soma, especially in the long-lived reproductives? And second, will we find evidence that the presumed non-canonical somatic function takes place outside the nucleus?

As reported here and previously (Koubová *et al*., 2021), *P. simplex TERT* is expressed in multiple isoforms defined by three alternative transcription start sites and alternative splicing events in exons 1 and 9 (Figure 1A, Supplementary Figure S3). Splicing of the first exon (*psTERT1* vs. *psTERT2*) may affect the leading sequence, which likely serves as a switch for targeting the translated protein to the nucleus and/or cytoplasm. Alternative splicing in the penultimate exon 9 determines the expression of the full-length isoform *TERT-A* containing all functional motifs of the C-terminal extension, or the truncated variant *TERT-B* lacking conserved motifs essential for full TERT activity in humans (Banik *et al*., 2002). We first confirmed all four possible combinations of these splicing events in *P. simplex* whole-body isolates by direct PCR amplification using isoform-specific primers (Supplementary Figure S1).

Then, by using DEXseq analysis of RNA sequencing data, we found a significant drop in the representation of the terminal part of exon 9 in reproductive organs from young and mature neotenic queens, indicating the preference for full-length splice variant *TERT-A* in the germline. This suggest that the TERT/telomerase reported in the somatic organs in part consists from the truncated *TERT-B* variant. We further verified this observation using RT-qPCR analyses with isoform-specific primers targeting both the N-terminal and C-terminal variants. All four variants gradually decreased in their transcript abundances from the embryonic stage through larval stages to the worker and soldier phenotypes. An interesting pattern has been observed in the relative ratios of the alternative variants. The full length *psTERT-A* clearly dominates over the truncated *psTERT-B* in the embryos compared to workers, and in gonads of male and female neotenics compared to the soma of neotenics and workers. Inter-caste comparison indicated that the ratio of *psTERT-A* to *psTERT-B* is highly significantly superior in the soma of neotenics compared to the soma of workers. Among the N-terminal isoforms, the ratio of the likely extranuclear *psTERT1* to the nuclear *psTERT2* was found higher in embryonic and larval immature phenotypes compared to workers and soldiers. Both *psTERT1* and *psTERT2* are significantly more expressed in gonads compared to the soma of neotenics and workers, and their ratio is highly significantly biased towards the extranuclear *psTERT1* in the soma of neotenics compared to their gonads and to the soma of workers.

Taken together, these observations on C- and N-terminal splice variants allow us to propose that the canonical telomere extension role is represented by the nuclear and full length variant *psTERT2-A*, as judged from the conspicuous relative enrichment of variants 2 and A in the gonads of both sexes. By contrast, inter-caste comparison suggests that the full-length *psTERT1-A* having the extranuclear signal, dominates over other variants in the somatic organs of kings and queens compared to workers, and may thus be the variant responsible for the previously reported telomerase upregulation in somatic tissues of reproductives, which prompted the present study. This variant is thus the candidate for the hypothesized non-canonical function independent of telomeres and nuclear localization, especially activated in kings and queens, even though present also in other castes and developmental stages. Identification of *psTERT1-A* as the variant upregulated in reproductives is further supported by the fact that the soma of kings and queens displays a greatly elevated reverse transcriptase activity manifested in TRAP (Telomerase Repeated Amplification Protocol) assays (Koubová *et al*., 2021), suggesting that the responsible TERT variant should have retained the telomerase activity, just as it seems likely for the full-length *psTERT1-A*.

The role of alternative splicing in the function and activity of TERT holoenzyme has been extensively studied during last decade in humans, but remains unclear in other species. In humans, the expression of *hTERT* mRNA is very low compared to other genes and only up to 5% of its transcripts correspond to full-length *TERT* isoform containing all 16 protein coding exons in both, the normal and cancer cell lines. The remaining transcripts represent other *hTERT* splice variants (Yi *et al*., 2001). Alternative splicing was shown to exert specific patterns in human development and thus serve as a post-transcriptional mechanism to tightly control the telomerase activity (Ulaner *et al*., 1998, 2001; Yi *et al*., 1999). Up to date, there are around 22 known spliced *hTERT* isoforms most of which are combination products of the major splice events listed in Supplementary Table S5. The most studied *hTERT* splice variants are minus-alpha, beta, gamma and delta4-13, which affect the reverse transcriptase domain (RT) encoded by exons 5–9, inhibit enzymatic activity and result in impaired telomere maintenance. All variants capable of binding the telomerase RNA component (TERC) are commonly considered as dominant-negative isoforms involved in post-transcriptional regulation of *hTERT* (Colgin *et al*., 2000; Yi *et al*., 2000, 2001; Penev *et al*., 2021). Another isoform identified in vertebrates and likely involved in post-transcriptional regulation is the DEL2 lacking the second exon. Skipping of exon 2 leads to frame shift, introduces preliminary termination codon in the subsequent exon and addresses the non-functional truncated transcript to nonsense-mediated decay (Hrdličková *et al*., 2012a; Withers *et al*., 2012). Some alternatives of DEL2 retaining partial sequence of exon 2 while keeping all other exons in frame were found in platypus and chicken (Hrdličková *et al*., 2012b), but even those are unlikely to participate in telomeres maintenance due to deletion of domains essential for proper binding of the RNA component.

Nevertheless, none of the listed variants affecting either the TERT active site or TERC binding domain was detected in *P. simplex.* On the other hand, the effect of alternative splicing in exon 9 giving rise to *psTERT-B* resembles to *hTERT* variants INS3 and INS4 (Saebøe-Larssen *et al*., 2006). All three variants introduce a preliminary termination codon in similar position of the C-terminal extension resulting in truncated protein lacking conserved motifs EIII and EIV. Deletion of these regions, especially the EIII conserved motif, leads to hTERT inactivation and loss of reverse transcriptase activity in the TRAP assay (Banik *et al*., 2002; Huard *et al*., 2003). Both, the INS3 and INS4 human splice isoforms were detected primarily in telomerase-positive cells and associated with heavy polysomes, thus suggesting a potential role for these variants in post-translational regulation of telomerase by acting in a dominant-negative fashion (Zhu *et al*., 2014). Our comparison in *P. simplex* showed that the ratio between *psTERT-B* and the full-length A variants was significantly higher in the soma of workers compared to soma and gonads of reproductive; this would confirm the enzymatic inactivity of *psTERT-B* as judged from low somatic telomerase activity in workers (Koubová *et al*., 2021).

To the best of our knowledge, the alternative *TERT* splicing has not been functionally studied in insects, even though the rapidly growing pool of publicly available insect sequences reveals the existence of diverse splicing patterns in various insect orders. Cross-taxa comparison among multiple species of Hymenoptera and Hemiptera, published in Lai et al. (2017), shows that in several included species *TERT* undergoes the splicing affecting the N-terminal region, giving rise to isoforms with unchanged reading frame comparable to *psTERT2*. Likewise, alternative splicing in the penultimate exon, characteristic of *psTERT-B*, and resulting in the truncated C-terminal extension was observed in several species (Lai *et al*., 2017). In light of the similarities in the telomerase activation in the long-lived reproductive phenotypes between termites and the honeybee, the latter should be targeted by future research with respect to the expression pattern of its *TERT* splicing variants in different castes and organs.

We addressed the question of subcellular distribution of somatic TERT both by ELISA protein quantification in cell compartments and using microscopic techniques. Our observations indicate that an important portion of TERT in somatic organs is localized outside the nucleus. Unlike in workers, the extranuclear TERT in abdomen of reproductives is more abundant than that found in nuclei, and its ratio to the nuclear TERT is significantly higher in neotenic kings and queens of two age classes compared to workers. Even though the anti-psTERT antibodies cannot retrieve individual TERT isoforms, it is reasonable to speculate that the extranuclear TERT corresponds to the *psTERT1-A* transcript variant. Comparison of abdominal psTERT abundance in nucleus, cytosol and mitochondria revealed its presence in all three compartments and suggests that it is specifically enriched in the cytosolic fraction of reproductives compared to workers. At the same time, the psTERT isoform pattern in the cytosol differs from that observed in the nucleus and mitochondria. Likewise, the immunodetection of psTERT in sections from various *P. simplex* organs revealed an abundant anti-psTERT signal in the cytoplasm of somatic tissues, including the ventral nerve cord and fat body (typical by low or no cell division incidence), even though we were not able to reliability compare the quantitative patterns among workers and reproductives.

All above observations point at the extranuclear localization of a specific TERT isoform in somatic tissues of different termite castes, which is particularly enriched in the soma of long-lived reproductives and may be linked with their longevity via an unknown non-canonical function independent of telomere maintenance. Within the growing list of non-canonical roles attributed to hTERT and telomerase in humans, several functions are accompanied by active telomerase export from the nucleus upon oxidative stress and participation of TERT in protection against oxidative damage (reviewed, e.g., in Cong & Shay, 2008; Ding *et al*., 2013; Saretzki, 2014; Romaniuk *et al*., 2019; Ségal-Bendirdjian & Geli, 2019; Zheng *et al*., 2019; Smith *et al*., 2022; Denham, 2023). It is mostly manifested in mitochondria by reducing the levels of reactive oxygen species, protecting mtDNA from oxidative damage and enhancing the efficiency of respiratory chain. These capacities are sometimes accompanied and/or dependent on reverse transcriptase function of telomerase (e.g., Fu *et al*., 2000; Ahmed *et al*., 2008; Haendeler *et al*., 2009; Indran *et al*., 2011). Telomerase shuttle to mitochondria is mediated via the N-terminal mitochondrial targeting sequence MTS and is phosphorylation-dependent (Liu *et al*., 2001; Haendeler *et al*., 2003; Santos *et al*., 2004). In other words, under certain physiological conditions, the same isoform acting in the nucleus is transferred to mitochondria to exert different roles. Our results indicate that such a mechanism is conserved also in termites as we observed a non-negligible psTERT abundance in the mitochondrial protein fraction from abdominal soma of queens and workers and the isoform profiles in both mitochondria and nucleus were similar.

Antioxidative action of telomerase/TERT in somatic mitochondria of termite kings and queens would be a plausible explanation of telomerase recruitment in reproductives given the variety of other antioxidant mechanisms the reproductives invest in. However, our results point at cytosolic rather than mitochondrial enrichment of extranuclear TERT in kings and queens, and the cytosolic isoform appears clearly different from the nuclear and mitochondrial pattern. Compared to body of knowledge on hTERT roles in human mitochondria, its potential functions in cytosol are understudied. Some studies report interaction of hTERT with several signaling pathways related to inflammation (NF-kappaB), but especially to metabolism and physiology. For instance, hTERT was shown to participate in regulation of glucose uptake in an insulin insensitive manner (Shaheen *et al*., 2014; Wardi *et al*., 2014). In vertebrates, cytosolic localization of TERT was repeatedly reported in multiple cell types of brain, especially in neurons which do not necessarily require telomere maintenance. Telomerase was shown to protect rodent embryonic hippocampal neurons from apoptosis during the development but to be subsequently downregulated in fully developed cells (Fu *et al*., 2000, 2002). Other studies revealed that the TERT mRNA and protein are present also in adults and their expression corelates more with pathological conditions than with age (Klapper *et al*., 2001; Spilsbury *et al*., 2015). Another study focusing on TERT localization in rat hippocampal neurons reported that neuronal aging is accompanied by the translocation of TERT from the nucleus to the cytoplasm. Only cytoplasmic TERT was observed in fully differentiated neurons both *in vitro* and *in situ* where the majority associated with nucleoprotein stress granules and participated in regulation of the cyclin kinase inhibitor p15INK4B (Iannilli *et al*., 2013). We did not observe such structures in termite neural tissue, but, surprisingly, we repeatedly observed psTERT localized to protein granular particles in fat body cells.

## Conclusions

In the present study, we confirm our previous postulates (Koubová *et al*., 2021) on differential pattern of TERT expression and protein abundance in the soma of sterile workers and long-lived kings and queens of *P. simplex*. We show that this caste-specific pattern is determined by differential expression of several *psTERT* splice variants, some of which are preferentially expressed in the germline, while the others in the somatic cells, including postmitotic organs, and that the ratios of psTERT somatic isoforms differs between workers and reproductives. Because the psTERT isoforms differ in their localization signals, also the resulting abundance of psTERT in somatic cell compartments differs among castes. In particular, we highlight the enrichment of the full-length psTERT1-A isoform in the extranuclear (more specifically cytosolic) fraction of abdominal soma of kings and queens compared to workers. By contrast, the nuclear and mitochondrial somatic fractions appear similar in the isoforms present and in their abundance in different castes. As judged from its protein sequence, the psTERT1-A should have retained its reverse transcriptase function and may thus account for the increased telomerase activity in the soma of reproductives, documented previously in TRAP assays (Koubová *et al*., 2021). It would be of particular interest to study the *TERT* splicing, its caste and tissue specificity and pattern of subcellular distribution in workers and queens of the honeybee, another representative of social insects with high telomerase activities in postmitotic somatic organs (Korandová & Čapková Frydrychová, 2016). This could confirm the convergent evolution of telomerase/TERT recruitment for non-canonical pro-longevity action in the soma of long-lived reproducing castes of advanced social insects. Our observations also allow for two general conclusions about the biological significance of telomerase. First, telomerase/TERT in insects behaves as a pleiotropic enzyme whose roles seem to encompass more functions than the canonical telomere extension in the germline, embryonic and stem cells, as evidenced by its expression in various somatic organs independently of their cell replication potential and its presence in various cell compartments. This resonates with the current view of telomerase as a protein adopting various roles according to the life histories of different metazoan lineages, which go far beyond the telomere maintenance. And second, our results demonstrate once again that the complex alternative splicing of *TERT* may be the key to the mentioned pleiotropy, allowing its tight developmental regulation in the canonical function as well as its manifestation in the non-canonical, sometimes lineage-specific functions.

## Supporting information

Electronic supplementary material

## Acknowledgements

The study was supported by the Charles University Grant Agency (project GA UK No 338021) and the Institute of Organic Chemistry and Biochemistry, CAS (RVO: 61388963).

## Author contribution

O. L. and R. Han. designed the study, J. K. prepared all termite samples, M. P. performed all isolations and both, the molecular biology and immunochemistry analyses, M. V. and H. S. performed immunohistochemistry analyses and confocal fluorescent microscopy, R. Had. was responsible for immunogold labeling and transmission electron microscopy imaging, O. L. performed NGS and statistical data analyses, O. L., R. Han and M. P. drafted the manuscript. All authors contributed to the text of the manuscript and approved its final version.

## Data availability

Raw data from RNA-seq experiments used for comparison of *psTERT* expression and differential exon usage in multiple *P. simplex* castes and body parts are published in NCBI SRA database under accession numbers XYXYXYX (*to be completed upon publication*).

