## Supplementary material for "Extended longevity of termite kings and queens is accompanied by extranuclear localization of telomerase in somatic organs and caste-specific expression of its isoforms": Electronic supplementary material

**Table S1.** List of primers used for qPCR-based expression profiling of *psTERT* isoforms

| Primer name | Target region | Orientation | Sequence | Product length | Primer efficiency |
|---|---|---|---|---|---|
| psTERT-altTSS_A-F | TERT1 | F | CGCTGTTACAGAGCTGTGGTC | 182 bp | 102.5 % |
| psTERT-altTSS_A-R | | R | TCAGTGCTGACATTTGGCTTTA | | |
| psTERT-TSS1-qPCR-F | TERT2 | F | TCCAACGTGTAATTGTTCAGACA | 148 bp | 101.3 % |
| psTERT_EX2_R1 | | R | ACACAAGACTTGGTTGAAGATGGA | | |
| psTERT-TES-qPCR-F2 | TERT-B | F | AGATATTGCCCATTCCATGC | 340 bp | 105.7 % |
| psTERT-TES1-qPCR-R | | R | TCTGTCTCTATATTTTCTTGGCGC | | |
| psTERT-TES-qPCR-F2 | TERT-A | F | AGATATTGCCCATTCCATGC | 351 bp | 99.9 % |
| psTERT-TES2-qPCR-R3 | | R | TACATCATCCAGCGAACAGC | | |
| psGAPDH-F | GAPDH | F | GTCGCTTCAAGGGTGAAGTT | 166 bp | 98.3 % |
| psGAPDH-R | | R | GATGCCTTTTCGATGGTTGT | | |

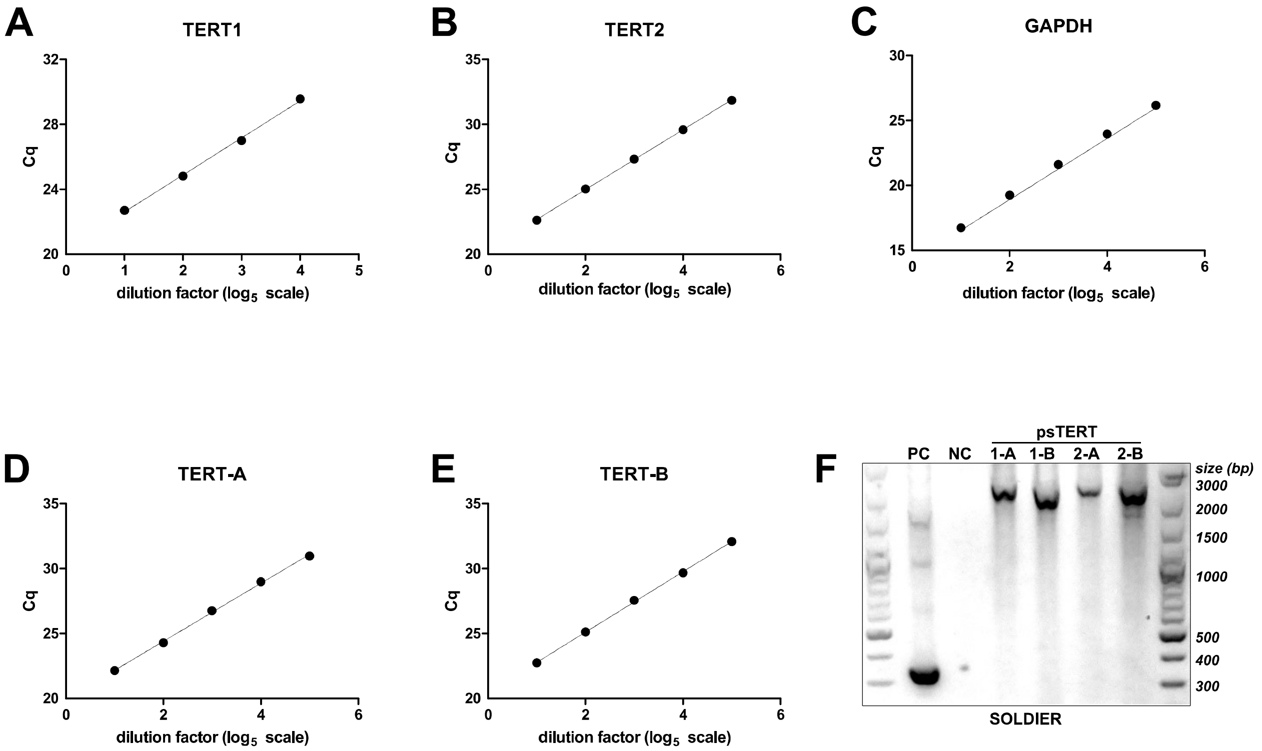


**Figure S1.** PCR detection of *psTERT* isoforms. **A-E**. Calibration curves of Cq values for cDNA serial dilution series used for calculation of qPCR primer efficiencies of *TERT1, TERT2, TERT-A, TERT-B* and *GAPDH* assays. **F**. PCR detection of all four splice variant combinations *TERT1-A, TERT1-B, TERT2-A* and *TERT2-B* in cDNA isolated from *P. simplex* soldiers. PC - positive control prepared using primers specific for splice variant *psTERT1*, NC - no template control.

**Table S2.** Expression of *psTERT* in RNA-seq data from various castes and tissues

| Caste | Tissue | Sample ID | Total number of reads | TERT read counts | FPKM | FPKM mean | FPKM SD |
|---|---|---|---|---|---|---|---|
| mature neotenic gueen (>2 years old) | abdominal soma | MFB1 | 5,248,186 | 78 | 5.48 | **5.64** | *0.48* |
| | | MFB2 | 5,046,248 | 71 | 5.19 | | |
| | | MFB3 | 7,122,462 | 122 | 6.32 | | |
| | | MFB4 | 7,643,969 | 115 | 5.55 | | |
| mature neotenic gueen (>2 years old) | reproductive organs | MFG1 | 9,277,670 | 256 | 10.18 | **12.58** | *2.16* |
| | | MFG2 | 9,366,589 | 296 | 11.66 | | |
| | | MFG3 | 7,322,612 | 302 | 15.22 | | |
| | | MFG4 | 6,321,494 | 227 | 13.25 | | |
| mature neotenic gueen (>2 years old) | head | MFH1 | 5,551,562 | 160 | 10.63 | **10.84** | *0.15* |
| | | MFH2 | 7,986,297 | 238 | 11.00 | | |
| | | MFH3 | 4,880,298 | 144 | 10.89 | | |
| | | MFH4 | 8,680,936 | 255 | 10.84 | | |
| pseudergate (worker) | abdominal soma | WB1 | 7,966,238 | 130 | 6.02 | **6.36** | *0.47* |
| | | WB2 | 8,509,544 | 143 | 6.20 | | |
| | | WB3 | 8,099,659 | 155 | 7.06 | | |
| | | WB4 | 8,334,976 | 139 | 6.15 | | |
| pseudergate (worker) | head | WH3 | 7,108,756 | 235 | 12.20 | **11.69** | *0.71* |
| | | WH4 | 9,899,490 | 294 | 10.96 | | |
| | | WH5 | 5,370,084 | 163 | 11.20 | | |
| | | WH6 | 6,403,463 | 215 | 12.39 | | |
| young neotenic gueen (3< month old) | abdominal soma | YFB1 | 7,457,596 | 128 | 6.33 | **7.50** | *1.57* |
| | | YFB2 | 6,351,011 | 116 | 6.74 | | |
| | | YFB3 | 7,457,854 | 198 | 9.80 | | |
| | | YFB4 | 6,983,818 | 135 | 7.13 | | |
| young neotenic gueen (3< month old) | reproductive organs | YFG1 | 8,760,328 | 412 | 17.35 | **15.32** | *2.24* |
| | | YFG2 | 9,471,560 | 366 | 14.26 | | |
| | | YFG3 | 6,377,926 | 294 | 17.01 | | |
| | | YFG4 | 7,278,176 | 250 | 12.68 | | |
| young neotenic gueen (3< month old) | head | YFH1 | 8,560,172 | 154 | 6.64 | **11.05** | *4.27* |
| | | YFH2 | 9,626,068 | 296 | 11.35 | | |
| | | YFH6 | 6,938,810 | 285 | 15.16 | | |

**Two-way ANOVA for abdominal soma and heads of workers, young queens and old queens**

| Source of Variation | % of total variation | F | P value |
|---|---|---|---|
| Interaction | 0.38 | 0.08582 | 0.9182 |
| Body part Factor | 54.02 | 24.27 | 0.0001 |
| Caste Factor | 9.46 | 2.126 | 0.1500 |

**t-test for gonads in young vs. mature queens**

| t | P value | df |
|---|---|---|
| 1.496 | 0.1852 | 6 |

**Table S3a.** Test statistics relative to the results presented in Figure 2A–F

**TERT-A (Figure 2A)**

| BROWN-FORSYTHE test | | | | |
|---|---|---|---|---|
| Df: | 6, 21 | | | |
| F: | 1.1762 | | | |
| P value: | 0.3561 | | | |
| **ONE-WAY ANOVA** | | | | |
| Df: | 6, 21 | | | |
| F: | 67.6236 | | | |
| P value: | <0.0001 | | | |
| **DUNNETT'S POST HOC TEST** | | | | |
| | n | Mean Diff. | q | Summary |
| W | 4 | | | |
| E | 4 | 2.1812 | 14.28 | *** |
| L1 | 4 | 0.3575 | 2.340 | ns |
| L2 | 4 | 0.3237 | 2.119 | ns |
| L3 | 4 | 0.0487 | 0.3190 | ns |
| preS | 4 | -0.2575 | 1.686 | ns |
| S | 4 | -0.5525 | 3.617 | ** |

**TERT-B (Figure 2B)**

| BROWN-FORSYTHE test | | | | |
|---|---|---|---|---|
| Df: | 6, 21 | | | |
| F: | 2.5720 | | | |
| P value: | 0.052 | | | |
| **ONE-WAY ANOVA** | | | | |
| Df: | 6, 21 | | | |
| F: | 34.2769 | | | |
| P value: | <0.0001 | | | |
| **DUNNETT'S POST HOC TEST** | | | | |
| | n | Mean Diff. | q | Summary |
| W | 4 | | | |
| E | 4 | 1.7662 | 11.34 | *** |
| L1 | 4 | 0.4749 | 3.050 | * |
| L2 | 4 | 0.5537 | 3.556 | ** |
| L3 | 4 | 0.3287 | 2.111 | ns |
| preS | 4 | 0.0125 | 0.08031 | ns |
| S | 4 | -0.1363 | 0.8755 | ns |

**TERT-B / TERT-A (Figure 2C)**

| BROWN-FORSYTHE test | | | | |
|---|---|---|---|---|
| Df: | 6, 21 | | | |
| F: | 0.9412 | | | |
| P value: | 0.4870 | | | |
| **ONE-WAY ANOVA** | | | | |
| Df: | 6, 21 | | | |
| F: | 9.6250 | | | |
| P value: | <0.0001 | | | |
| **DUNNETT'S POST HOC TEST** | | | | |
| | n | Mean Diff. | q | Summary |
| W | 4 | | | |
| E | 4 | -0.4150 | 3.331 | * |
| L1 | 4 | 0.1175 | 0.9434 | ns |
| L2 | 4 | 0.2300 | 1.846 | ns |
| L3 | 4 | 0.2800 | 2.248 | ns |
| preS | 4 | 0.2700 | 2.168 | ns |
| S | 4 | 0.4163 | 3.342 | * |

**TERT-A (Figure 2D)**

| BROWN-FORSYTHE test | | | |
|---|---|---|---|
| Df: | 4, 14 | | |
| F: | 0.4317 | | |
| P value: | 0.7835 | | |
| **ONE-WAY ANOVA** | | | |
| Df: | 4, 14 | | |
| F: | 220.6 | | |
| P value: | <0.0001 | | |
| **POST HOC (Bonferroni mult. comp.)** | | | |
| | Mean Diff. | t | Summary |
| W / NQ | -0.08000 | 0.5851 | ns |
| W / NK | -0.1225 | 0.8498 | ns |
| NQ / NK | -0.04249 | 0.3107 | ns |
| NQ / O | -3.420 | 22.97 | *** |
| NK / T | -2.361 | 15.16 | *** |

**TERT-B (Figure 2E)**

| BROWN-FORSYTHE test | | | |
|---|---|---|---|
| Df: | 4, 14 | | |
| F: | 2.916 | | |
| P value: | 0.0601 | | |
| **ONE-WAY ANOVA** | | | |
| Df: | 4, 14 | | |
| F: | 5.304 | | |
| P value: | 0.0082 | | |
| **POST HOC (Bonferroni mult. comp.)** | | | |
| | Mean Diff. | t | Summary |
| W / NQ | 0.3581 | 3.278 | * |
| W / NK | 0.3887 | 3.376 | * |
| NQ / NK | 0.03062 | 0.2803 | ns |
| NQ / O | -0.3033 | 2.551 | ns |
| NK / T | -0.3449 | 2.774 | ns |

**TERT-B / TERT-A (Figure 2F)**

| BROWN-FORSYTHE test | | | |
|---|---|---|---|
| Df: | 4, 14 | | |
| F: | 2.4171 | | |
| P value: | 0.0978 | | |
| **ONE-WAY ANOVA** | | | |
| Df: | 4, 14 | | |
| F: | 43.14 | | |
| P value: | <0.0001 | | |
| **POST HOC (Bonferroni mult. comp.)** | | | |
| | Mean Diff. | t | Summary |
| W / NQ | 1.058 | 5.41 | *** |
| W / NK | 1.228 | 5.958 | *** |
| NQ / NK | 0.1702 | 0.8698 | ns |
| NQ / O | 1.482 | 6.961 | *** |
| NK / T | 1.068 | 4.797 | ** |

*Signif. codes:* ***, $p < 0.001$; **, $p < 0.01$; *, $p < 0.05$

*Experimental groups:*

*E ... egg*
*L1 - 3 ... larval stages*
*W … worker*
*preS ... presoldier*
*S ... soldier*
*NQ ... neotenic queen*
*NK ... neotenic king*
*O ... ovaria*
*T ... testes*

**Table S3b.** Test statistics relative to the results presented in Figure 2G–L

| TERT1 (Figure 2G) | | | | |
|---|---|---|---|---|
| BROWN-FORSYTHE test | | | | |
| Df: | 6, 21 | | | |
| F: | 0.4501 | | | |
| P value: | 0.8367 | | | |
| ONE-WAY ANOVA | | | | |
| Df: | 6, 21 | | | |
| F: | 134.1 | | | |
| P value: | <0.0001 | | | |
| DUNNETT'S POST HOC TEST | | | | |
| | n | Mean Diff. | q | Summary |
| W | 4 | | | |
| E | 4 | -3.836 | 23.10 | *** |
| L1 | 4 | -0.7912 | 4.765 | *** |
| L2 | 4 | -0.8587 | 5.172 | *** |
| L3 | 4 | -0.4887 | 2.943 | * |
| preS | 4 | -0.2025 | 1.220 | ns |
| S | 4 | 0.1013 | 0.6099 | ns |

| TERT2 (Figure 2H) | | | | |
|---|---|---|---|---|
| BROWN-FORSYTHE test | | | | |
| Df: | 6, 21 | | | |
| F: | 1.375 | | | |
| P value: | 0.2702 | | | |
| ONE-WAY ANOVA | | | | |
| Df: | 6, 21 | | | |
| F: | 15.68 | | | |
| P value: | <0.0001 | | | |
| DUNNETT'S POST HOC TEST | | | | |
| | n | Mean Diff. | q | Summary |
| W | 4 | | | |
| E | 4 | -1.418 | 6.402 | *** |
| L1 | 4 | -0.06250 | 0.2823 | ns |
| L2 | 4 | -0.1275 | 0.5759 | ns |
| L3 | 4 | 0.2050 | 0.9258 | ns |
| preS | 4 | 0.5387 | 2.433 | ns |
| S | 4 | 0.1387 | 0.6263 | ns |

| TERT1 / TERT2 (Figure 2I) | | | | |
|---|---|---|---|---|
| BROWN-FORSYTHE test | | | | |
| Df: | 6, 21 | | | |
| F: | 1.109 | | | |
| P value: | 0.3900 | | | |
| ONE-WAY ANOVA | | | | |
| Df: | 6, 20 | | | |
| F: | 15.59 | | | |
| P value: | <0.0001 | | | |
| DUNNETT'S POST HOC TEST | | | | |
| | n | Mean Diff. | q | Summary |
| W | 4 | | | |
| E | 4 | -0.4319 | 4.101 | ** |
| L1 | 4 | -0.6584 | 6.252 | *** |
| L2 | 4 | -0.6608 | 6.275 | *** |
| L3 | 4 | -0.6191 | 5.878 | *** |
| preS | 4 | -0.6756 | 6.415 | *** |
| S | 3 | -0.06425 | 0.6101 | ns |

| TERT1 (Figure 2J) | | | |
|---|---|---|---|
| BROWN-FORSYTHE test | | | |
| Df: | 4, 15 | | |
| F: | 0.2404 | | |
| P value: | 0.9110 | | |
| ONE-WAY ANOVA | | | |
| Df: | 4, 15 | | |
| F: | 32.55 | | |
| P value: | <0.0001 | | |
| POST HOC (Bonferroni mult. comp.) | | | |
| | Mean Diff. | t | Summary |
| W / NQ | -0.6825 | 2.460 | ns |
| W / NK | -0.2975 | 1.221 | ns |
| NQ / NK | 0.3850 | 1.451 | ns |
| NQ / O | -1.139 | 4.104 | ** |
| NK / T | -2.093 | 8.586 | *** |

| TERT2 (Figure 2K) | | | |
|---|---|---|---|
| BROWN-FORSYTHE test | | | |
| Df: | 4, 15 | | |
| F: | 0.1430 | | |
| P value: | 0.9633 | | |
| ONE-WAY ANOVA | | | |
| Df: | 4, 15 | | |
| F: | 99.38 | | |
| P value: | <0.0001 | | |
| POST HOC (Bonferroni mult. comp.) | | | |
| | Mean Diff. | t | Summary |
| W / NQ | 0.08088 | 0.3499 | ns |
| W / NK | 0.4515 | 2.224 | ns |
| NQ / NK | 0.3706 | 1.677 | ns |
| NQ / O | -2.966 | 12.83 | *** |
| NK / T | -2.516 | 12.39 | *** |

| TERT1 / TERT2 (Figure 2L) | | | |
|---|---|---|---|
| BROWN-FORSYTHE test | | | |
| Df: | 4, 15 | | |
| F: | 0.4390 | | |
| P value: | 0.9980 | | |
| ONE-WAY ANOVA | | | |
| Df: | 4, 15 | | |
| F: | 24.69 | | |
| P value: | <0.0001 | | |
| POST HOC (Bonferroni mult. comp.) | | | |
| | Mean Diff. | t | Summary |
| W / NQ | -0.7663 | 4.585 | ** |
| W / NK | -0.7202 | 4.907 | *** |
| NQ / NK | 0.04601 | 0.2879 | ns |
| NQ / O | 1.318 | 7.887 | *** |
| NK / T | 0.4358 | 2.969 | * |

*Signif. codes:* ****, p < 0.001; **, p < 0.01; *, p < 0.05*

*Experimental groups:*

*E ... egg*
*L1 - 3 ... larval stages*
*W … worker*
*preS ... presoldier*
*S ... soldier*
*NQ ... neotenic queen*
*NK ... neotenic king*
*O ... ovaria*
*T ... testes*

**Table S4.** Test statistics relative to the results presented in Figure 3

| ELISA somatic TERT (Figure 3A - extranuclear) | | | | |
|---|---|---|---|---|
| **BROWN-FORSYTHE test** | | | | |
| Df: | 4, 10 | | | |
| F: | 0.2587 | | | |
| P value: | 0.8977 | | | |
| **ONE-WAY ANOVA** | | | | |
| Df: | 4, 10 | | | |
| F: | 7.039 | | | |
| P value: | 0.0058 | | | |
| **DUNNETT'S POST HOC TEST** | | | | |
| | n | Mean Diff. | q | Summary |
| W | 3 | | | |
| YQ | 3 | -1.170 | 3.675 | * |
| YK | 3 | -1.474 | 4.630 | ** |
| MQ | 3 | -1.271 | 3.993 | ** |
| MK | 3 | -0.6227 | 1.956 | ns |

| ELISA somatic TERT (Figure 3A - nuclear) | | | | |
|---|---|---|---|---|
| **BROWN-FORSYTHE test** | | | | |
| Df: | 4, 10 | | | |
| F: | 0.538 | | | |
| P value: | 0.712 | | | |
| **ONE-WAY ANOVA** | | | | |
| Df: | 4, 10 | | | |
| F: | 1.645 | | | |
| P value: | 0.2383 | | | |
| **DUNNETT'S POST HOC TEST** | | | | |
| | n | Mean Diff. | q | Summary |
| W | 3 | | | |
| YQ | 3 | -0.003509 | 0.01248 | ns |
| YK | 3 | 0.4741 | 1.686 | ns |
| MQ | 3 | 0.3826 | 1.361 | ns |
| MK | 3 | 0.5108 | 1.817 | ns |

| ELISA somatic TERT extranuclear/nuclear (Figure 3B) | | | | |
|---|---|---|---|---|
| **BROWN-FORSYTHE test** | | | | |
| Df: | 4, 10 | | | |
| F: | 0.2209 | | | |
| P value: | 0.9207 | | | |
| **ONE-WAY ANOVA** | | | | |
| Df: | 4, 10 | | | |
| F: | 10.39 | | | |
| P value: | 0.0014 | | | |
| **DUNNETT'S POST HOC TEST** | | | | |
| | n | Mean Diff. | q | Summary |
| W | 3 | | | |
| YQ | 3 | -1.166 | 3.577 | * |
| YK | 3 | -1.948 | 5.974 | *** |
| MQ | 3 | -1.654 | 5.072 | ** |
| MK | 3 | -1.133 | 3.476 | * |

| ELISA TERT in cytoplasm, nucleus , mitochondria (Figure 3C) | | | | | |
|---|---|---|---|---|---|
| cytoplasm | | nucleus | | mitochondria | |
| **F test for variances** | | **F test for variances** | | **F test for variances** | |
| Df: | 2, 3 | Df: | 3, 2 | Df: | 2, 2 |
| F: | 3.355 | F: | 3.186 | F: | 8.165 |
| P value: | 0.3434 | P value: | 0.4982 | P value: | 0.2182 |
| **t-test** | | **t-test** | | **t-test** | |
| Df: | 5 | Df: | 5 | Df: | 4 |
| t: | 2.604 | t: | 0.2594 | t: | 0.117 |
| P value: | 0.0474 | P value: | 0.8057 | P value: | 0.9125 |

| ELISA extranuclear TERT in gonads (Figure 3E) | | | | |
|---|---|---|---|---|
| **BROWN-FORSYTHE test** | | | | |
| Df: | 4, 11 | | | |
| F: | 0.7890 | | | |
| P value: | 0.5560 | | | |
| **ONE-WAY ANOVA** | | | | |
| Df: | 4, 11 | | | |
| F: | 2.261 | | | |
| P value: | 0.1284 | | | |
| **DUNNETT'S POST HOC TEST** | | | | |
| | n | Mean Diff. | q | Summary |
| W | 3 | | | |
| YQ | 3 | 0.1859 | 1.051 | ns |
| YK | 3 | 0.3825 | 2.163 | ns |
| MQ | 3 | 0.1937 | 1.171 | ns |
| MK | 3 | 0.4806 | 2.718 | ns |

*Signif. codes:* ****, p < 0.001; **, p < 0.01; *, p < 0.05*

*Experimental groups:*

*W ... worker*
*YQ ... young queen (<6 months)*
*MQ ... mature queen (>2 years)*
*YK ... young king (<6 months)*
*MK ... mature king (>2 years)*

```
                                                                                   psTERT-altTSS_A-F
psTERT gene     1 AATTGCTAACGAAAACCAACAAAATGCACAAATGTTTCGGTCGTGCTTGACCATTTTCAGGGTGCGCTGTTACAGAGTAAGTGATACAACGACGTGTTCA 100
psTERT1           AATTGCTAACGAAAACCAACAAAATGCACAAATGTTTCGGTCGTGCTTGACCATTTTCAGGGTGCGCTGTTACAGA------------------------
                                                  M  F  R  S  C  L  T  I  F  R  V  R  C  Y  R  -  -  -  -  -  -  -  -
                                                          putative signal peptide
psTERT2β          AATTGCTAACGAAAACCAACAAAATGCACAAATGTTTCGGTCGTGCTTGACCATTTTCAGGGTGCGCTGTTACAGAGTAAGTGATACAACGACGTGTTCA
psTERT2                                                                  TTCAGGGTGCGCTGTTACAGAGTAAGTGATACAACGACGTGTTCA

                        psTERT-TSS1-qPCR-F                            exon 1 ┐┌ exon 2
psTERT gene   101 TTCGTCCAACGTGTAATTGTTCAGACATATATTTCCACAATCCAAATATCGTGTGGCTGTGGTCTTCTGGGTTGTGACACTGTGTAGTCTTTTGACAGA 200
psTERT1           --------------------------------------------------------GCTGTGGTCTTCTGGGTTGTGACACTGTGTAGTCTTTTGACAGA
                   -  -  -  -  -  -  -  -  -  -  -  -  -  -  -  -  -  -  -  A  V  V  F  W  V  V  T  L  C  S  L  L  T  D

psTERT2           TTCGTCCAACGTGTAATTGTTCAGACATATATTTCCACAATCCAAATATCGTGTGGCTGTGGTCTTCTGGGTTGTGACACTGTGTAGTCTTTTGACAGA

                                                  psTERT_EX2_R1 exon 2 ┐┌ exon 3
psTERT gene   201 CTGCTGTTGTTTTGAGAAAGCTACTCTATCCATCTTCAACCAAGTCTTGTGTTGGATGAACAAAGACATTTCTCACAAGCATATAAGAAGAAGAAAAAAG 300
psTERT1           CTGCTGTTGTTTTGAGAAAGCTACTCTATCCATCTTCAACCAAGTCTTGTGTTGGATGAACAAAGACATTTCTCACAAGCATATAAGAAGAAGAAAAAAG
                    C  C  C  F  E  K  A  T  L  S  I  F  N  Q  V  L  C  W  M  N  K  D  I  S  H  K  H  I  R  R  R  K  K

psTERT2           CTGCTGTTGTTTTGAGAAAGCTACTCTATCCATCTTCAACCAAGTCTTGTGTTGGATGAACAAAGACATTTCTCACAAGCATATAAGAAGAAGAAAAAAG
                                                                          M  N  K  D  I  S  H  K  H  I  R  R  R  K  K
                                                                                                       nuclear signal
                        psTERT-altTSS_A-R
psTERT gene   301 AAAATAAAGCCAAATGTCAGCACTGACAGCAGAATAGGATACAAGGGAACAACCGTATTGTCAAACAAACAAATGCTGTATAACCAAGCTACACATGAGC 400
psTERT1           AAAATAAAGCCAAATGTCAGCACTGACAGCAGAATAGGATACAAGGGAACAACCGTATTGTCAAACAAACAAATGCTGTATAACCAAGCTACACATGAGC
                   K  I  K  P  N  V  S  T  D  S  R  I  G  Y  K  G  T  T  V  L  S  N  K  Q  M  L  Y  N  Q  A  T  H  E

psTERT2           AAAATAAAGCCAAATGTCAGCACTGACAGCAGAATAGGATACAAGGGAACAACCGTATTGTCAAACAAACAAATGCTGTATAACCAAGCTACACATGAGC
                   K  I  K  P  N  V  S  T  D  S  R  I  G  Y  K  G  T  T  V  L  S  N  K  Q  M  L  Y  N  Q  A  T  H  E

                                                                       ...

                                            psTERT-TES-qPCR-F2                       exon 8 ┐┌ exon 9
psTERT gene  2001 TGATTATGCTCGCTACAGGTTTCAAGATATTGCCCATTCCATGCGTCTGACAGAAGTGAAATGTCCTGGAAAGTACCTTCAGAAGCGGATGGAAGGTTCA 2100
psTERT-A,B        TGATTATGCTCGCTACAGGTTTCAAGATATTGCCCATTCCATGCGTCTGACAGAAGTGAAATGTCCTGGAAAGTACCTTCAGAAGCGGATGGAAGGTTCA
                    D  Y  A  R  Y  R  F  Q  D  I  A  H  S  M  R  L  T  E  V  K  C  P  G  K  Y  L  Q  K  R  M  E  G  S


psTERT gene  2101 TTTTCTTCATTAAAACTGAATGTTCTTGTCCTGGATGAGCTGTACAACTCAAAAGATGTAGTTCTCACAAACGTATTTTATGCAGGTCTTTTGCAGGCTT 2200
psTERT-A,B        TTTTCTTCATTAAAACTGAATGTTCTTGTCCTGGATGAGCTGTACAACTCAAAAGATGTAGTTCTCACAAACGTATTTTATGCAGGTCTTTTGCAGGCTT
                  F  S  S  L  K  L  N  V  L  V  L  D  E  L  Y  N  S  K  D  V  V  L  T  N  V  F  Y  A  G  L  L  Q  A  F


psTERT gene  2201 TCAGATTCCACAGTCTTGTTAAATGTACCTTTGATAAGAAGAATCTGAATCAGCAGTTTCTTACCCAAATAGTGTATGAAACTGTCGGTAGGACTGGGGG 2300
psTERT-A,B        TCAGATTCCACAGTCTTGTTAAATGTACCTTTGATAAGAAGAATCTGAATCAGCAGTTTCTTACCCAAATAGTGTATGAAACTGTCGGTAGGACTGGGGG
                  R  F  H  S  L  V  K  C  T  F  D  K  K  N  L  N  Q  Q  F  L  T  Q  I  V  Y  E  T  V  G  R  T  G  G

                                                              psTERT-TES1-qPCR-R             exon 9 ┐┌ exon 10
psTERT gene  2301 ACACACATTGAATATTGCCAAGAAATACTGCAAAGGTAATGCGCCAAGAAAATATAGAGACAGAATAAACCTCTATTAATCCACTTCGACGCTTGACTTA 2400
psTERT-B          ACACACATTGAATATTGCCAAGAAATACTGCAAAGGTAATGCGCCAAGAAAATATAGAGACAGAATAAACCTCTATTAATCCA
                    H  T  L  N  I  A  K  K  Y  C  K  G  N  A  P  R  K  Y  R  D  R  I  N  L  Y  *

psTERT-A          ACACACATTGAATATTGCCAAGAAATACTGCAAAG------------------------------------------------CTTCGACGCTTGACTTA
                    H  T  L  N  I  A  K  K  Y  C  K  -  -  -  -  -  -  -  -  -  -  -  -  -  -  -  -  A  S  T  L  D  L
                     psTERT-TES2-qPCR-R3
psTERT gene  2401 AATGCTGTTCGCTGGATGATGTACAGAGCTTTCTTCTTGAGATTGTCCATTGTTCAAAGCTACTATCGTCTTGTACTGAGAAACATTGATTGGATGTTAA 2500
psTERT-A          AATGCTGTTCGCTGGATGATGTACAGAGCTTTCTTCTTGAGATTGTCCATTGTTCAAAGCTACTATCGTCTTGTACTGAGAAACATTGATTGGATGTTAA
                   N  A  V  R  W  M  M  Y  R  A  F  F  L  R  L  S  I  V  Q  S  Y  Y  R  L  V  L  R  N  I  D  W  M  L
                                                   conserved region E-III

psTERT gene  2501 AGCGTCTTTCTGAAAGGATACCAGATGAAAAATGTGATCAGATGATGCAGCTAACGTTACGCATTCCAGATGCATTTAAAACAGTTTCATAAAATGTTGC 2600
psTERT-A          AGCGTCTTTCTGAAAGGATACCAGATGAAAAATGTGATCAGATGATGCAGCTAACGTTACGCATTCCAGATGCATTTAAAACAGTTTCATAAAATGTTGC
                  K  R  L  S  E  R  I  P  D  E  K  C  D  Q  M  M  Q  L  T  L  R  I  P  D  A  F  K  T  V  S  *
                                                                  conserved region E-IV

psTERT gene  2601 CGCGCTTCTGCATGAACATGTTAGGTACTGGTGTGCCATGTGACACAGTCTTTAGGCACTTAACTTTGATTGCTTAATATGAACAAATAGTTGGCTAGAT 2700
psTERT-A          CGCGCTTCTGCATGAACATGTTAGGTACTGGTGTGCCATGTGACACAGTCTTTAGGCACTTAACTTTGATTGCTTAATATGAACAAATAGTTGGCTAGAT


psTERT gene  2701 GACCAAGCC                                                                                            2709
psTERT-A          GACCAAGCC
```

**Figure S2. Nucleotide and protein sequence alignment representing alternative splicing in exons 1 and 9 and in isoforms TERT1, 2 and TERT-A, B.** Genomic reference is in bold and shaded in gray, primer annealing sites used for isoform-specific expression analyses are underlined and shown in blue (forward) and green (reverse). Functionally important regions such as putative signal peptide (pink), nuclear signal (orange) and C-terminal conserved regions EIII and EIV (both blue) are shaded and bold-faced. Numbers represent positions in full transcript sequence containing all *psTERT* exons, exon junctions are represented by broken lines.

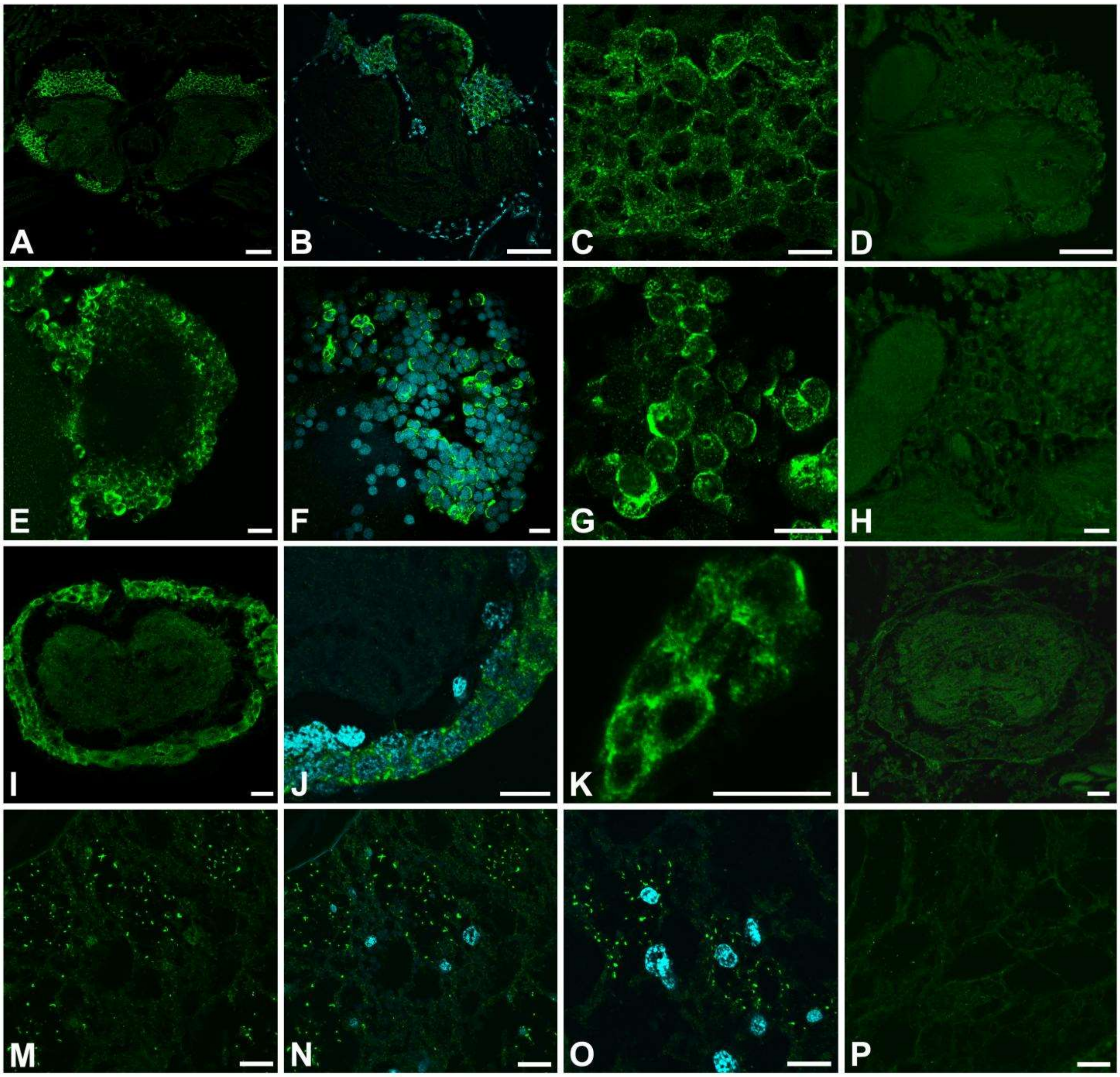


**Figure S3. Immunodetection of psTERT in the central nervous system and in the fat body of *P. simplex* workers**. **A-C.** Paraplast sections with psTERT positive staining in brain cells. **D.** Absence of positive signal in the control brain section when primary antibody was replaced by preimmune serum. **E-G.** Immunodetection of psTERT in wholemount preparations of the brain revealed an identical staining pattern as on paraplast sections. The images show different magnifications of the optic lobe region. **H.** High magnification image of the control sample shown in D. **I-K.** Different magnifications of psTERT immunoreactivity in paraplast section of an abdominal ganglion of the ventral nerve cord. **L.** Absence of positive signal in a ganglion of the control sample. **M-O.** Different magnifications of psTERT immunoreactivity in paraplast sections of fat body. **P.** Absence of positive signal in the control fat body sample. In B, F, J, N and O, the psTERT signal (green) is merged with DAPI staining (blue). Scale bars represent 50 µm in A, B, D and 10 µm in C, E-P.

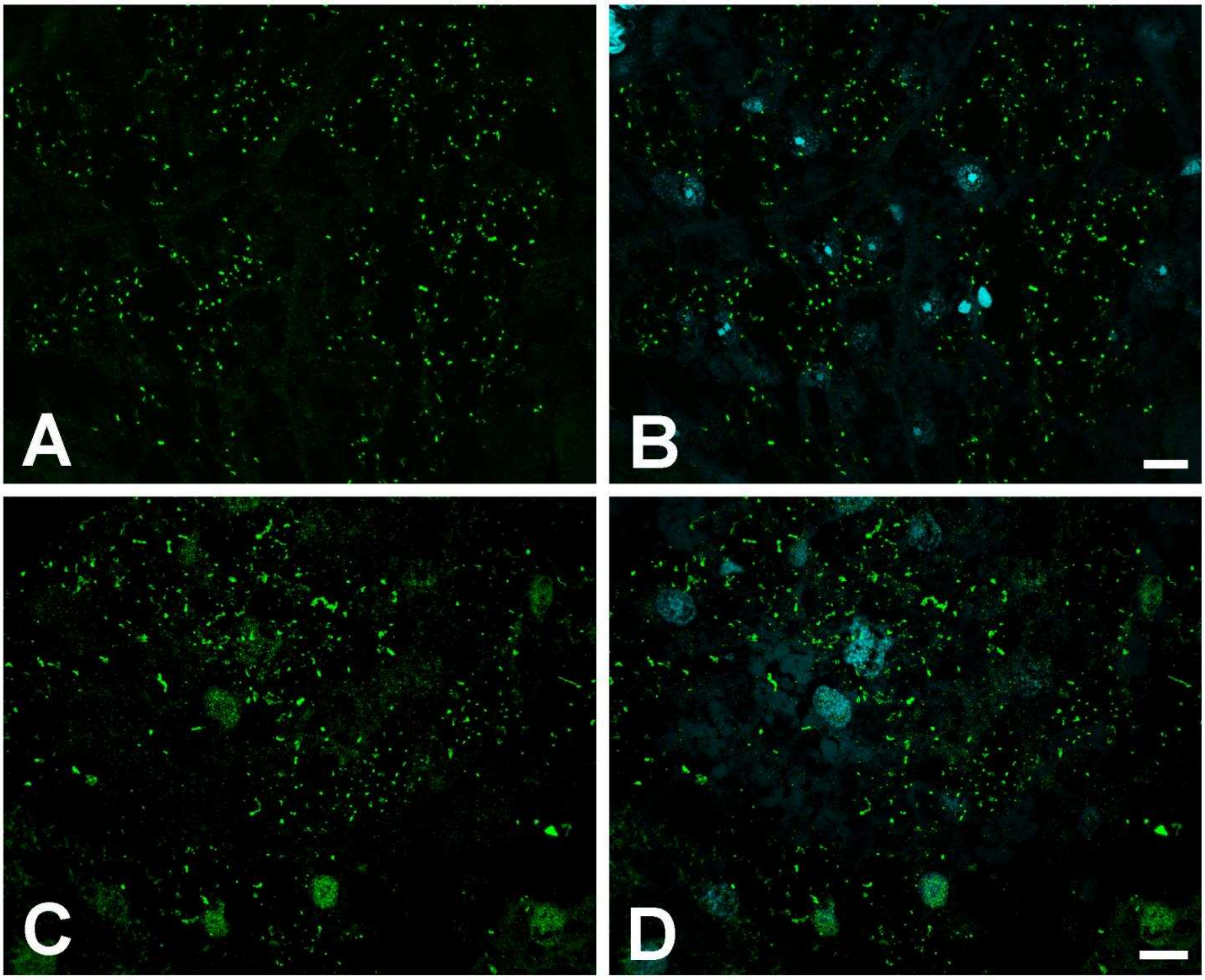


**Figure S4. Abundant psTERT-RP immunoreactivity in fat body of some queens and kings. A, B.** Fat body of queen. **C, D.** Fat body of king. Left - labelling with anti-psTERT-RP antibody, right - psTERT-RP signal (green) merged with DAPI staining (blue). Scale bar = 10 μm.

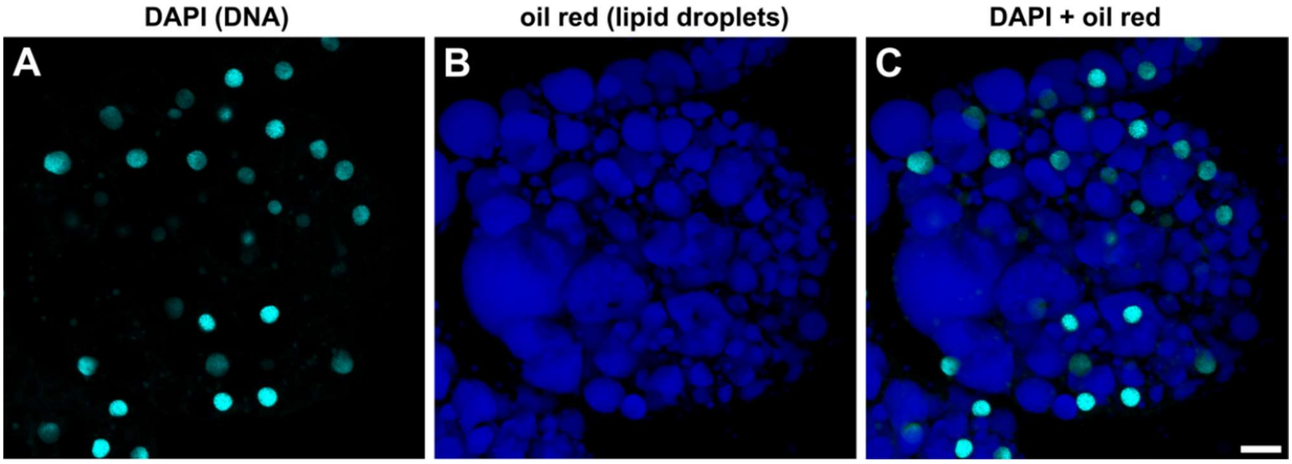


**Figure S5. Structure of adipous tissue of the fat body in workers.** Paraplast section of the fat body stained with DAPI (A), oil red (B) and DAPI + oil red (C) to visualize lipid droplets and nuclei in adipocytes. The extranuclear cytoplasm is marginalized by the large lipid droplets, filling the majority of the cell volume. Scale bar represents 15 μm.

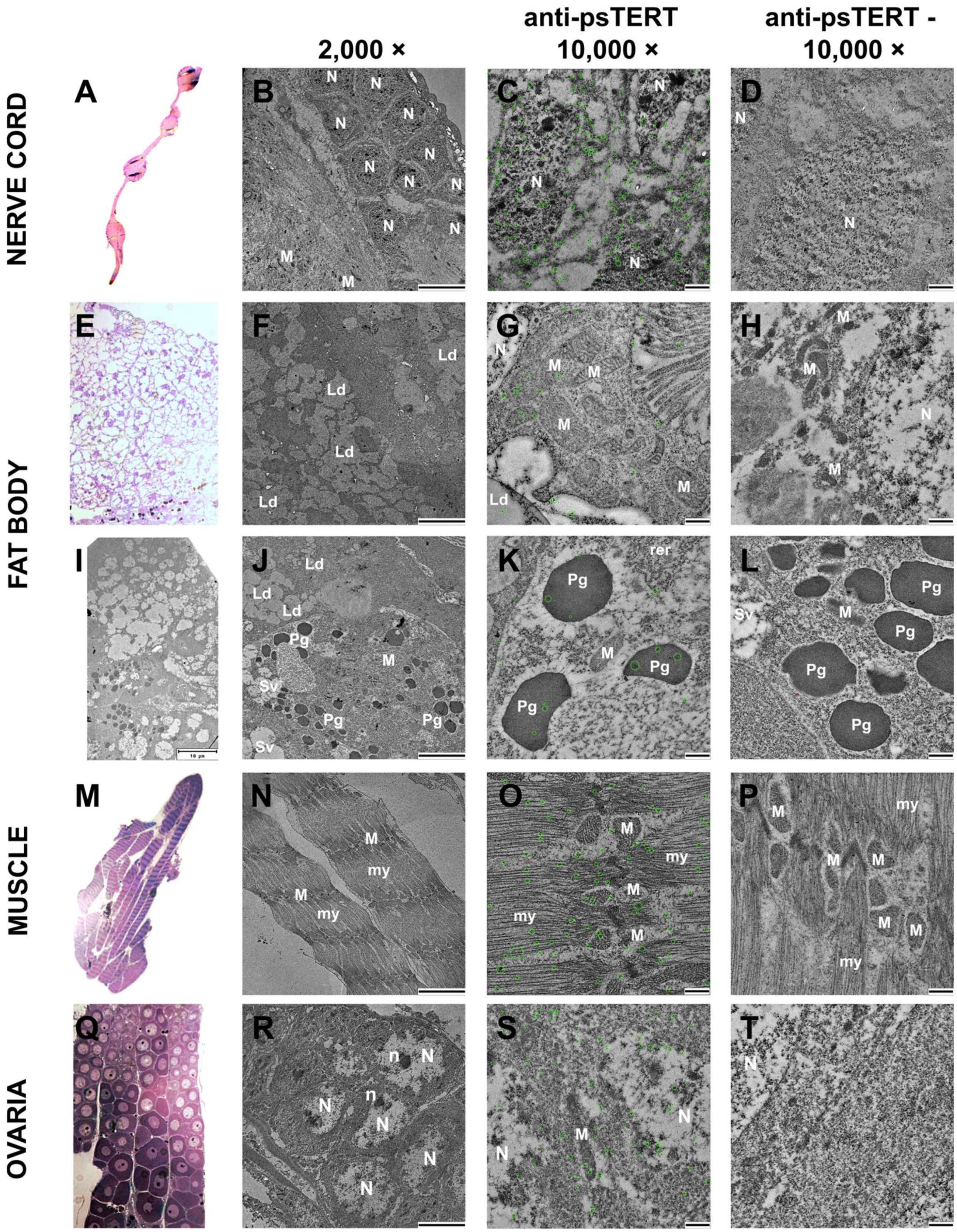
2,000 ×
anti-psTERT 10,000 ×
anti-psTERT - 10,000 ×
NERVE CORD
FAT BODY
MUSCLE
OVARIA
A
B
C
D
E
F
G
H
I
J
K
L
M
N
O
P
Q
R
S
T
N
M
Ld
Pg
Sv
rer
my
n

**Figure S6. Transmission electron microscopy (TEM) with psTERT detection using anti-psTERT antibody and immunogold labeling.** Immunoreactions were performed on ultrathin sections of tissue samples dissected from *P. simplex* neotenic queens: ventral nerve cord (A-B), abdominal fat body (E-L), muscle (M-P) and ovaria (Q-T). **(A, E, M, Q)**. Semithin sections of examined tissues stained with toluidine blue and taken under optical microscope. **I.** TEM micrograph at 1000x magnification showing multiple cell types in the fat body tissue. **(B,F,J,N,R)**. TEM micrographs at 2000x magnification showing structure of examined tissues, scale bar 5 µm. **(C,G,K,O,S)**. TEM micrographs at 10000x magnification with immunochemical detection of psTERT, scale bar 500 nm. Cellular organelles are marked with white symbols: N - nucleus, n - nucleolus, M -mitochondria, Ld - lipid droplets, Sv - secretory vesicles, Pg -protein granules, rer - rough endoplasmic reticulum, my - myofibrils.

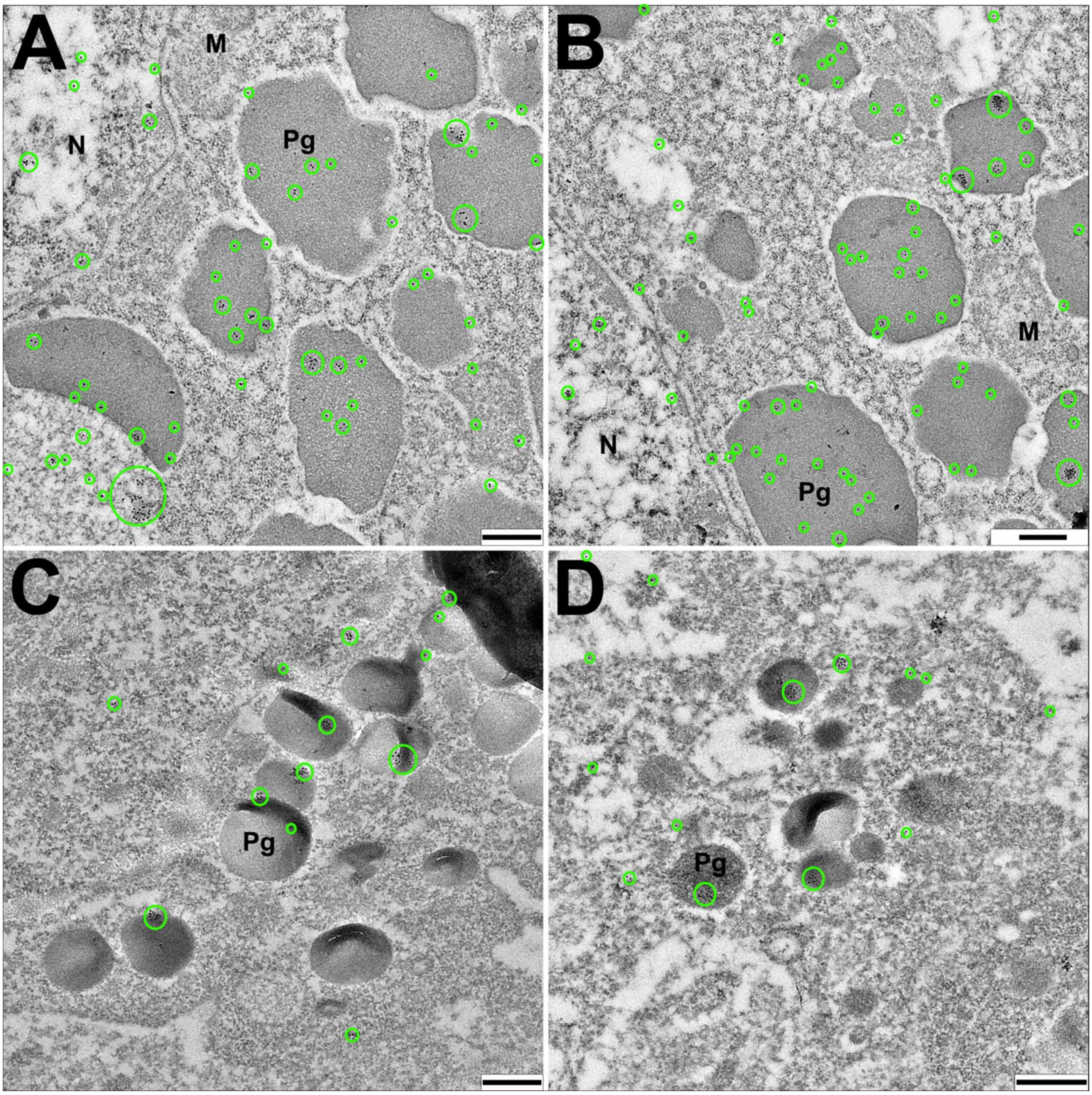


**Figure S7. Transmission electron microscopy (TEM) with immunogold detection showing psTERT accumulation to two types of protein granules on ultrathin sections of fat body from *P. simplex* neotenic queens. (A, B)**. Typical proptein granules from adipous tissue of *P. simplex* abdominal fat body. **(C, D)**. Protein granules on fat body sections found commonly in the proximity of other organs such as reproductive organs or ventral nerve cord. Cellular organelles are marked by black symbols: N - nucleus, M - mitochondria, Pg - protein granules. Scale bar represents 500 nm.

**Table S5.** Most frequent splice events observed in hTERT according to Ludlow *et al.* (2019).

| isoform | Splicing event | Termination codon | RT activity | Function |
|---|---|---|---|---|
| **full-length** | none | Standard, exon 16 | full | Telomere maintenance |
| **DEL2** | Deletion of exon 2 | PTC, exon 3 | No | Mostly degraded |
| **minus-alpha** | Partial deletion of exon 6 | Standard, exon 16 | No | Dominant-negative, binds *hTERC* |
| **minus-beta** | Deletion of exons 7, 8 | PTC, exon 10 | No | Mostly degraded, may play a role in DNA repair |
| **minus gamma** | Deletion of exon 11 | Standard, exon 16 | No | Dominant-negative, binds hTERC, tissue-specific |
| **delta4-13** | Deletion of exons 4-13 | Standard, exon 16 | No | Proposed to stimulate proliferation |
| **INS3** | Partial insertion of intron 14 | PTC, intron 14 | partial | Dominant-negative, binds *hTERC*, tissue-specific |
| **INS4** | Partial insertion of intron 14 | PTC, exon 14 | partial | Dominant-negative, binds *hTERC*, tissue-specific |

*Legend: PTC - preliminary termination codon, hTERC – human telomerase RNA component*
